# RIM and RIM-binding protein localize synaptic CaV2 channels to differentially regulate transmission in neuronal circuits

**DOI:** 10.1101/2021.02.01.429206

**Authors:** Barbara Jánosi, Jana F. Liewald, Szi-chieh Yu, Simon Umbach, Ivan C. Alcantara, Amelie C.F. Bergs, Martin Schneider, Jiajie Shao, Alexander Gottschalk

**Author notes:** these authors contributed equally (Barbara Jánosi, Jana F. Liewald).

## Abstract

At chemical synapses, voltage-gated Ca^2+^-channels (VGCCs) translate electrical signals into a trigger for synaptic vesicle (SV) fusion. VGCCs and the Ca^2+^ microdomains they elicit must be located precisely to primed SVs, to evoke rapid transmitter release. Localization is mediated by Rab3 interacting molecule (RIM) and RIM-binding proteins (RIM-BPs), which interact and bind to the C-terminus of the CaV2 VGCC α-subunit. We studied this machinery at the mixed cholinergic/GABAergic neuromuscular junction (NMJ) of *Caenorhabditis elegans. rimb-1* mutants had mild synaptic defects, through loosening the anchoring of UNC-2/CaV2 and delaying the onset of SV fusion. UNC-10/RIM deletion much more severely affected transmission. Even though postsynaptic depolarization was reduced, *rimb-1* mutants had increased cholinergic (but reduced GABAergic) transmission, to compensate for the delayed release. This did not occur when the excitation-inhibition balance was altered by removing GABA transmission. RIMB-1 thus may differentially regulate transmission in mixed circuits. Untethering the UNC-2/CaV2 channel by removing its C-terminal PDZ ligand exacerbated the *rimb-1* defects, and similar phenotypes resulted from acute degradation of the CaV2 β-subunit CCB-1. Therefore, untethering of the CaV2 complex is as severe as its elimination, yet does not abolish transmission, likely due to compensation by CaV1. Thus, robustness and flexibility of synaptic transmission emerges from VGCC regulation.

## Introduction

Neurons release chemical signals by fusion of synaptic vesicles (SVs) with the plasma membrane (Brunger et al., 2018; Sudhof, 2013). This process occurs in synaptic terminals, triggered by action potentials that activate voltage gated Ca^2+^ channels (VGCCs), which mediate a rapid, local rise in cytosolic Ca^2+^ (Nanou and Catterall, 2018). This is detected by the Ca^2+^ sensor for synchronous release, synaptotagmin, an intrinsic component of SVs (Sudhof, 2012). Protein complexes intricately organize SVs and P/Q-type VGCCs (CaV2.1 in mammals, UNC-2 in *Caenorhabditis elegans*) in the presynaptic active zone membrane, to ensure an optimal distance for rapid Ca^2+^ sensing (Acuna et al., 2015; Acuna et al., 2016; Han et al., 2011; Held et al., 2020; Hibino et al., 2002; Kaeser et al., 2011; Liu et al., 2011; Wu et al., 2019). While SNARE complexes hold the SV in a configuration ready to catalyze membrane fusion in response to the Ca^2+^ signal, the VGCC, via its C-terminal tail, is held in place by a complex of the RIM (RAB3 interacting molecule) and RIM-binding proteins (RIM-BP). RIM binds to the SV via RAB3 proteins, and to other components of the active zone protein scaffold, termed presynaptic particle web in mammals, cytomatrix of the active zone (CAZ) in *Drosophila,* or dense projection (DP) in *C. elegans* (Ackermann et al., 2015).

In recent years the function of RIMBPs (RIMB-1 in *C. elegans*) was elucidated in mammalian and *Drosophila* neurons (Acuna et al., 2015; Acuna et al., 2016; Brockmann et al., 2019; Brockmann et al., 2020; Liu et al., 2011; Muller et al., 2015). While mice lacking both neuronal RIM-BPs show no alteration in number of hippocampal synapses and evoked release per se, the fidelity of precisely timed coupling of release in response to evoked action potentials is decreased (Acuna et al., 2015). RIM-BPs anchor CaV2 in the vicinity of primed SVs, by binding the VGCC C-terminus, however, they do not affect VGCC function or kinetics. The fly mutant *drbp* shows a more severe phenotype, with reduced Ca^2+^ sensitivity of release, and a defect in presynaptic homeostatic plasticity (Liu et al., 2011; Muller et al., 2015). This is because neurons cannot upregulate their Ca^2+^ influx, an otherwise normal response to the blocking of post-synaptic transmitter receptors (Muller and Davis, 2012). In *Drosophila, drbp* is genetically interacting with *rim* during baseline transmission, with similar phenotypes of both mutants, and more severe phenotypes if they are combined. RIM-BP is also affecting SV recycling after extensive synaptic activity (Muller et al., 2015). The synergy of RIM and RIM-BPs is also present in mice, ensuring the anchoring of active zone VGCCs, but affecting also the assembly of the active zone protein scaffold, as well as recruitment of Munc13 proteins, which are essential for SV priming (Acuna et al., 2016; Brockmann et al., 2019; Brockmann et al., 2020). However, in mice, RIM single mutants have a more pronounced defect in recruiting and stabilizing VGCCs at synapses than RIM-BP mutants (Han et al., 2011). In *C. elegans,* RIM deletion alone affects synaptic function more severely than in mice (Koushika et al., 2001). Transgenic overexpression of *C. elegans* GFP::UNC-2 (CaV2) protein can be augmented by overexpressing RIMB-1, and in the *rimb-1; unc-10* (RIM) double mutants, as in mice, synaptic abundance of CaV2/UNC-2 channels is reduced (Kushibiki et al., 2019). The molecular interplay of interactions of UNC-10 and RIMB-1 in locating the UNC-2 CaV2 channel has been analyzed in mutants lacking or overexpressing various proteins of the active zone protein scaffold (Oh et al., 2021). However, how RIMB-1 and RIM affect localization of UNC-2 relative to the docked SV is not known in *C. elegans,* and the synaptic ultrastructure of the *rimb-1* mutant is yet to be analyzed. Likewise, it is not clear how absence of RIMB-1 and the likely consequent untethering of UNC-2/CaV2 channels affects synaptic properties and the characteristics of chemical transmission in *C. elegans.*

The neuromuscular junction (NMJ) of *C. elegans* is thought to be driven by graded, not action potentials (Liu et al., 2009; Schultheis et al., 2011), though recent voltage imaging data suggest that cholinergic neurons may also produce fast, action-potential like activities (Azimi Hashemi et al., 2019). These result from activity of the premotor interneurons that are presynaptic to, and induce burst activity in, cholinergic motor neurons (Liu et al., 2013). The *C. elegans* NMJ is a tripartite synapse comprising cholinergic motor neurons that innervate muscle cells as well as GABAergic motor neurons. The latter project to the respective contralateral (dorso-ventral) side of the body to innervate opposing muscle. This circuitry orchestrates the dorso-ventral undulatory locomotion, and muscles thus integrate activity of both cholinergic and GABAergic inputs. Precise function of synapses in both types of neurons is required to ensure normal excitation-inhibition balance, and this likely involves precise positioning of VGCCs. In fact, cholinergic and GABAergic synapses exhibit a differential requirement of L- and P/Q-type VGCCs (Liu et al., 2018; Tong et al., 2017).

Here, we set out to characterize in detail the synaptic phenotypes of *rimb-1* mutants at the behavioral, physiological, and ultrastructural level, and to address molecular details of its interaction with UNC-2/CaV2 channels. Despite reduced postsynaptic depolarization, *rimb-1* mutants had increased, but delayed, cholinergic release, which occurred from SVs that were located distal to the center of the active zone. This homeostatic change was not found when inhibition at the mixed cholinergic/GABAergic NMJ was eliminated genetically. RIMB-1 interacts with RIM in localizing the CaV2 channel UNC-2 and we find that progressive de-anchoring of the UNC-2 channel, by C-terminal truncation and/or *rimb-1* mutation, causes phenotypes that resemble its deletion. RIMB-1 may differentially affect UNC-2 channels in different cell types, thus regulating the outcome of circuit function at different levels.

## Results

### RIMB-1 is expressed in cholinergic and GABAergic motor neurons

RIMB-1, the single *C. elegans* homolog of vertebrate RIM-BPs, contains three *src*-homology type III (SH3) domains, and three fibronectin type III (FN3) domains (**Fig. 1A**). RIM-BPs bind to proline-rich sequences of RIM (UNC-10) and the CaV2 α1 subunit (UNC-2) via their SH3 domains (Hibino et al., 2002; Wu et al., 2019). RIM/UNC-10 further binds to the C-terminal end of CaV2/UNC-2 via its PSD95/Discs large/Zonula occludens (PDZ) domain (Kaeser et al., 2011), thus the three proteins form a tripartite complex (**Fig. 1B**). To verify the expression of *rimb-1* in neurons, specifically at the NMJ, we co-expressed GFP (from the *rimb-1p* promoter, 3 kb upstream of the ATG) and mCherry (from the cholinergic neuron specific *unc-17p* promoter; **Fig. 1C**). *rimb-1p::GFP* expressed throughout the nervous system, including motor neurons in the ventral nerve cord, while *unc-17p*::mCherry was observed in a subset of neurons, in which it overlapped with *rimb-1p::GFP.* Thus, *rimb-1* is likely pan-neuronal, and at the NMJ, it is expressed in both cholinergic and GABAergic motor neurons.

**Figure 1:**
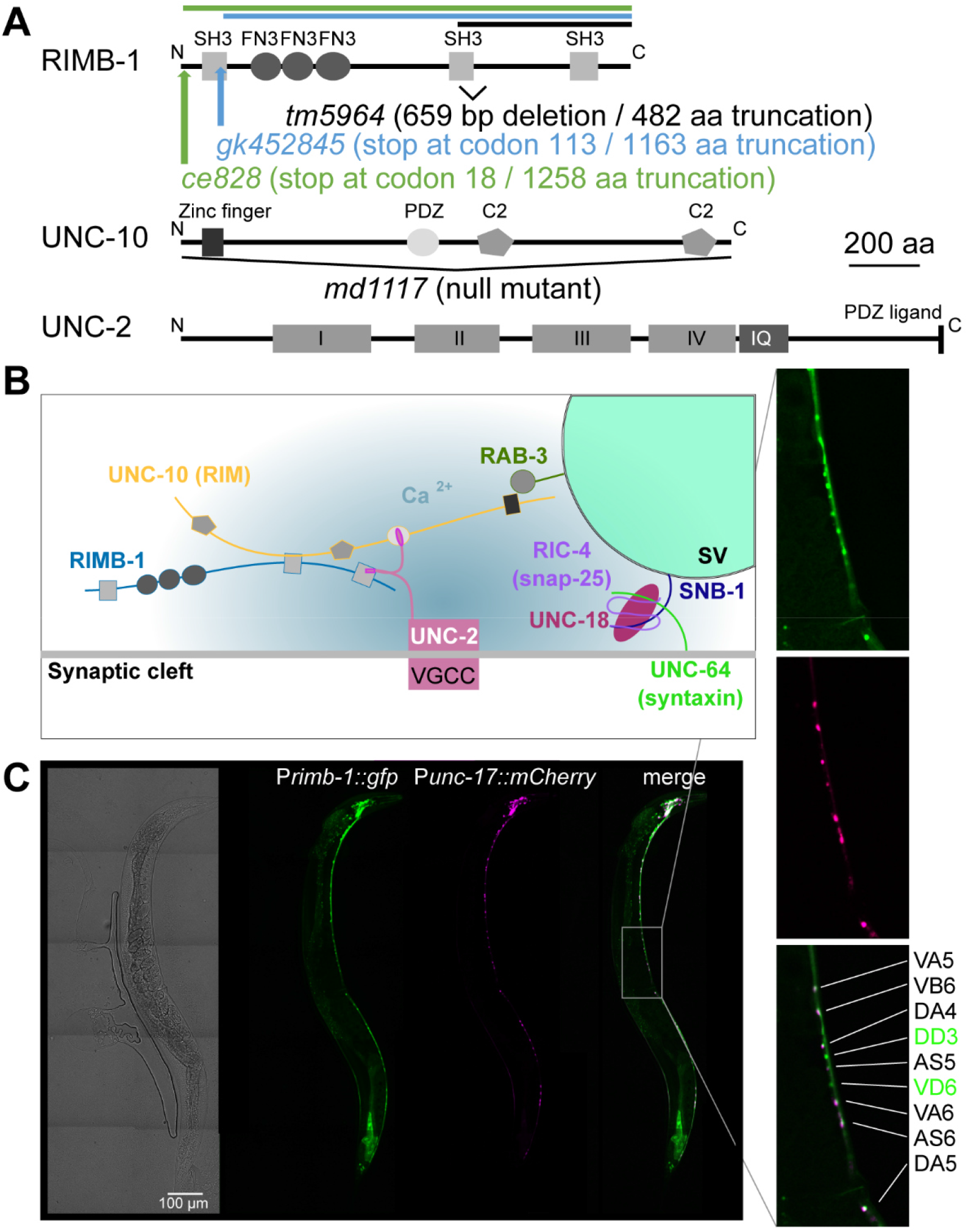
UNC-10/RIM and RIMB-1/RIM-binding protein affect UNC-2/CaV2 function in *C. elegans* (motor) neurons. **A)** Protein primary structures and sequence features (SH3 – src homology 3; FN3 – fibronectin 3; PDZ – PDZ domain; C2 – Ca^2+^ and phospholipid binding domain; I-IV – VGCC modules; IQ – IQ domain) of RIMB-1, UNC-10/RIM, and UNC-2/CaV2. Deletion sites / regions of alleles used in this work are indicated by bars or brackets. **B)** Arrangement of proteins anchoring the UNC-2 channel in the vicinity of a docked SV. **C)** Brightfield and fluorescence micrographs of an animal expressing GFP from the *rimb-1p* promoter and mCherry from the *unc-17p* promoter (cholinergic neurons). RIMB-1 appears to be pan-neuronal, and in the ventral nerve cord, is expressed in cholinergic (magenta and green) and GABAergic motor neurons (green only). Neuron designations: AS, DA, VA, VB, are cholinergic neurons required for locomotion (D, V: targeting dorsal or ventral partners); DD and VD are dorsally and ventrally innervating GABAergic motor neurons. Scale bar: 100 μm.

### *rimb-1* mutants exhibit reduced cholinergic transmission

We assessed possible defects in synaptic transmission of *rimb-1* animals. The *rimb-1(tm5964*) mutant has a 659 bp deletion affecting exon 14 and the preceding intron, which likely leads to a frameshift and early stop codon, resulting in a protein that misses the last 482 amino acids (**Fig. 1A**). Whether this represents a null allele is not known. In any case, this allele should affect the interaction of RIMB-1 and UNC-10/RIM, since it is mediated by the second SH3 domain (Wang et al., 2000), as well as the interaction with the UNC-2 VGCC, mediated by the third SH3 domain. Additional deletion alleles have been reported: *gk452845* truncates the protein already after amino acid 113, while *ce828* induces an early stop at codon 18 (**Fig. 1A**; Edwards et al., 2018). We compared the *tm5964* and *ce828* alleles. First we assayed cholinergic transmission: Incubation in the acetylcholinesterase inhibitor aldicarb leads to progressive paralysis due to accumulation of acetylcholine (ACh) in the synaptic cleft and muscle hyper-contraction (Mahoney et al., 2006). *rimb-1(tm5964*) animals paralyzed significantly later than wild type controls (**Fig. S1A**), while *ce828* animals essentially paralyzed with the same kinetics as wild type. To further compare these two alleles, we measured swimming and crawling behavior (**Fig. S1B, C**). Here, *ce828* animals were slightly, but significantly impaired, while *tm5964* animals showed a stronger defect. Thus, the *tm5964* allele, which likely retains the N-terminal two thirds of the protein, exhibits a stronger phenotype than the complete deletion of RIMB-1. Also a recent publication reported normal distribution of UNC-2 VGCCs in the *rimb-1(ce828*) mutant (Oh et al., 2021). Possibly, in the absence of RIMB-1, where also other interactions of its N-terminus with unknown partners are absent, and/or due to compensatory changes, the *ce828* animals show near wild type behavior. The *gk452845* allele, retaining 113 amino acids, had a locomotion defect and aldicarb resistance, similar to *tm5964* (Kushibiki et al., 2019). Thus, we used the *tm5964* allele in all experiments.

We compared *rimb-1(tm5964*) mutants to *unc-10(md1117*) animals, as well as *rimb-1; unc-10* double mutants, and wild type controls (**Fig. 2A**). Again, *rimb-1(tm5964*) animals paralyzed significantly later than controls on 2 mM aldicarb, indicating defective ACh release. *unc-10* mutants, as well as *rimb-1; unc-10* animals, were much more compromised, not paralyzing during the time course of the assay, thus emphasizing a very strong reduction of synaptic transmission. Function of post-synaptic nicotinic acetylcholine receptors (nAChRs) was assessed by a similar paralysis assay using an agonist of the main *C. elegans* muscle nAChR: The *rimb-1* mutant was similar to wild type in the levamisole sensitivity assay, in contrast to the nAChR subunit mutant *unc-38* (**Fig. S2A**). Thus, post-synaptic nAChRs are normal in *rimb-1* animals. Second, swimming behavior (**Fig. 2B**) was compromised in *rimb-1* mutants, which showed a significant reduction of the swimming rate, while *unc-10* as well as *rimb-1; unc-10* double mutants were almost paralyzed, with *rimb-1* exacerbating the *unc-10* single mutant phenotype. Third, we assessed cholinergic function by optogenetics. We expressed channelrhodopsin-2 (ChR2) in cholinergic neurons (transgene *zxIs6*) and photostimulated them for 60 s. This causes ACh release, muscle activation, and macroscopic body contraction (Liewald et al., 2008). Body length was measured before, during, and after photostimulation (**Fig. 2C, D**). Wild type animals contracted by about 10 %, *rimb-1* mutants only by about 6 %. *rimb-1; unc-10* double mutants initially shortened like wild type, but showed a significant, progressive reduction in the contraction to about 6 %. This is indicative of synaptic fatigue, possibly due to an SV recycling defect (Kittelmann et al., 2013). *rimb-1* mutants did not behave like typical SV release mutants (e.g., synaptobrevin *snb-1;* **Fig. S2B, C**), because such mutants show increased contraction, due to a compensatory or homeostatic increase in post-synaptic excitability (Liewald et al., 2008). When we directly excited muscles via ChR2 in *snb-1* and *rimb-1* mutants (**Fig. S2D**), *snb-1* mutants contracted more than controls, while *rimb-1* animals behaved like wild type. Thus, the abnormal contraction behavior of *rimb-1* mutants, evoked by ChR2 in cholinergic neurons, must be due to complex synaptic transmission defects that could affect not only SV release, but also recycling, possibly through VGCC signaling. As RIMB-1 is expressed in cholinergic and GABAergic motor neurons, it may affect NMJ transmission through both cell types.

**Figure 2:**
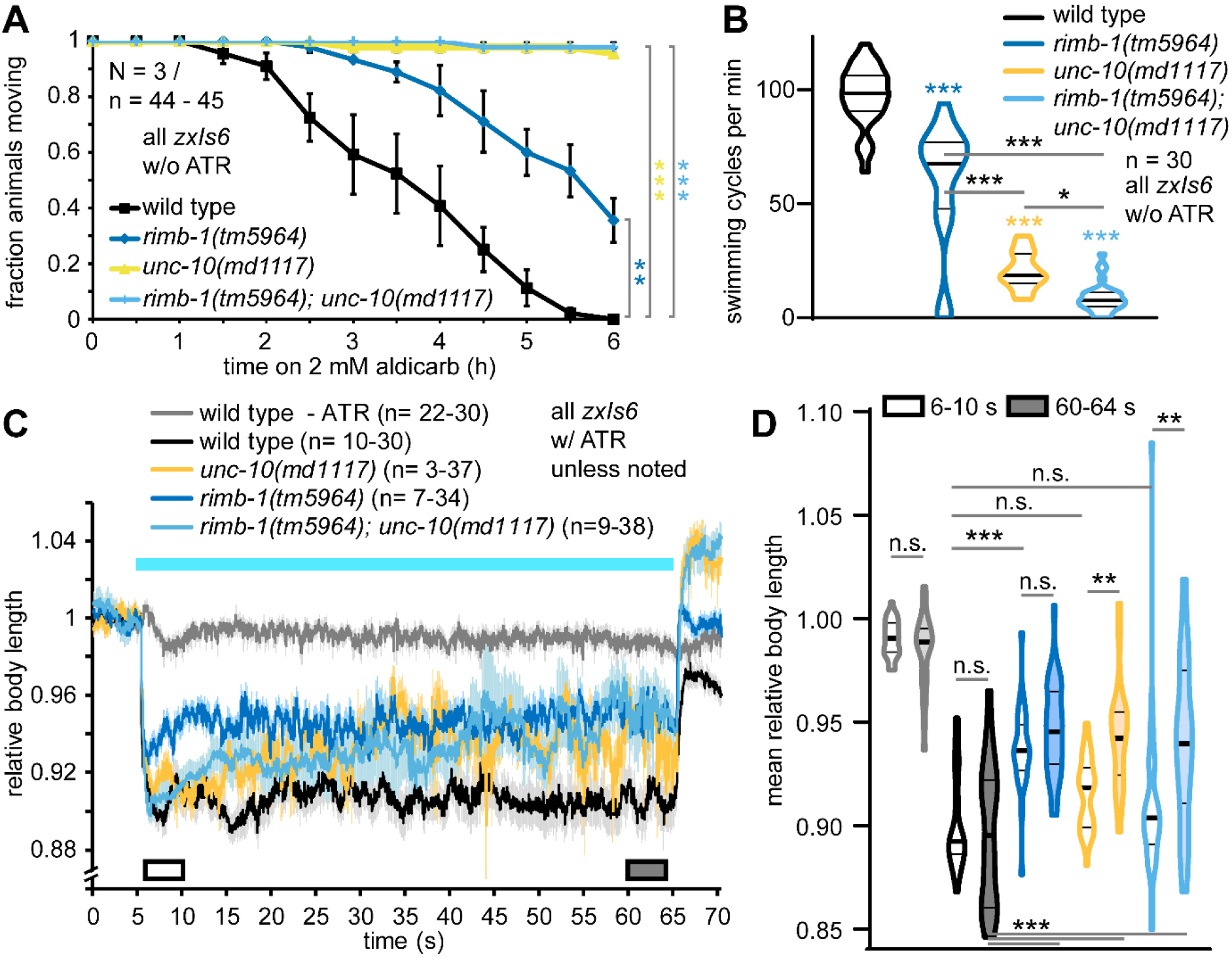
*rimb-1* mutants exhibit defective cholinergic transmission, locomotion and muscle activation. **A)** Aldicarb paralysis assay probing cholinergic transmission; progressive paralysis is delayed in mutants. The indicated genotypes and number of animals were scored and data from 3 experiments averaged at the indicated time points. Mean ± s.e.m. Log rank test, Bonferroni correction. **B)** Swimming locomotion assay of the indicated genotypes (n=30 animals each). Animals were left swimming in liquid and their full body thrashes during 1 min were counted. Data shown as median and 25/75 quartiles (thick and thin lines), min to max. Statistical significance determined by one-way ANOVA, Bonferroni-corrected. **C)** Photostimulated cholinergic transmission causes muscle contraction, measured by video microscopy. Relative body length (mean ± s.e.m.) of the indicated number of animals (note, for some time points, not all animals could be analyzed) of the indicated genotypes, before, during and after a 1 min photostimulus (blue bar). +/-ATR: presence/absence of all-*trans* retinal, rendering ChR2 functional (or not). White and grey boxes indicate periods for which statistical comparisons were calculated in **D)** Data in C) (6-10 and 60-64 s) shown as median and 25/75 quartiles (thick and thin lines), min to max. One-way ANOVA, Tukey test. In A, B, D: Statistical significance given as *p<0.05; **p<0.01; ***p<0.001.

### *rimb-1* mutants show delayed, but unexpectedly increased cholinergic ePSCs

To more directly assess transmission in *rimb-1(tm5964), unc-10(md1117),* and *rimb-1; unc-10* double mutants, we recorded post-synaptic currents in patch-clamped muscle cells. We analyzed spontaneous as well as evoked release, before and during repeated optogenetic stimulation *(via zxIs6;* **Fig. 3A**). *unc-10* and *rimb-1; unc-10* animals showed significantly reduced miniature post-synaptic current (mPSC) frequencies, while mPSC amplitudes were the same in all genetic backgrounds (**Fig. 3B, C**). Photostimulation evoked large PSCs (ePSCs); however, compared to wild type, ePSC amplitudes were unexpectedly increased in *rimb-1* mutants (first stimulus; **Fig. 3D, E**), and also remained larger throughout a stimulus train (0.5 Hz; 2-way ANOVA). In contrast, *unc-10* and *rimb-1; unc-10* animals showed significantly reduced ePSC amplitudes. The ePSC amplitude increase in *rimb-1* mutants was contradictory to the aldicarb and swimming assays as well as the evoked muscle contraction (**Fig. 2**), which were compromised in *rimb-1* mutants.

**Figure 3:**
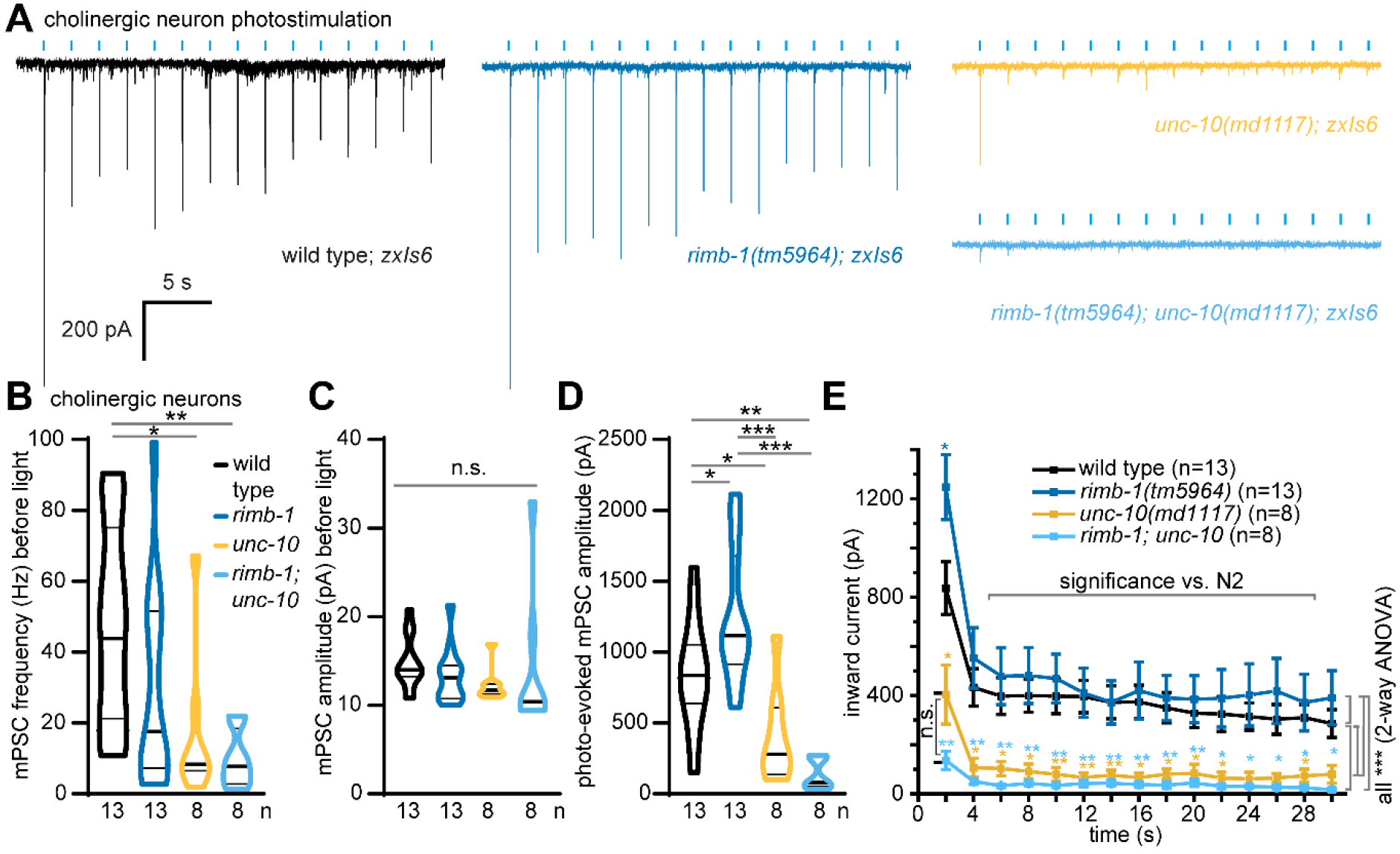
*rimb-1* and unc-10/RIM mutants exhibit increased and reduced cholinergic transmission at the NMJ. **A)** Photo-stimulation of cholinergic motor neurons induces post-synaptic currents. Shown are original records for the indicated genotypes, induced by 0.5 Hz, 10 ms stimuli (blue tick marks) of animals bearing transgene *zxIs6* (ChR2 in cholinergic neurons, expressed from promoter *unc-17p).* **B)** Miniature post-synaptic current (mPSC) frequencies and **C)** amplitudes during 30 s before first light stimulus. Data shown as median and 25/75 quartiles (thick and thin lines), min to max. One-way ANOVA, Kruskall-Wallis test. **D)** Photoevoked post-synaptic current (ePSC) amplitude of the first, and consecutive **E)** photostimuli (mean ± s.e.m.). One-way ANOVA with Tukey test in D, and for individual stimuli in E; two-way ANOVA in E to compare entire datasets. Statistical significance given as *p<0.05; **p<0.01; ***p<0.001.

### Delayed current onset determines reduced muscle depolarization in *rimb-1* mutants

Why are ePSCs increased in *rimb-1* mutants, despite the behavioral deficits? To explore possible reasons for this discrepancy, we analyzed parameters of evoked release: amplitude, delay from light pulse onset to peak current (time to peak), rise and decay times, as well as area under the curve (**Fig. 4A-D**). *rimb-1* mutants were delayed in their current onset and time to peak, particularly if combined with the *unc-10* RIM mutant. Quantal content, calculated by dividing the area under the ePSC curve by the mean area under the mPSC peak (**Fig. 4D**), was significantly smaller in the *rimb-1; unc-10* mutants. Both proteins thus contribute to effective synaptic transmission at the *C. elegans* NMJ, likely through jointly anchoring the UNC-2/CaV2 channel to SV release sites. We wondered if the timing of postsynaptic current onset may determine the outcome of muscle activation. *C. elegans* motor neurons produce bursts of transmitter release, which effect muscular action potentials (Liu et al., 2013). Timing of postsynaptic current events thus determines muscle activation and this is likely promoted by RIMB-1. We therefore analyzed the timing of photo-evoked release (**Fig. 4E, F**), looking at the first derivative (slope) of the current rise. These events were occurring 1-2 ms faster in wild type than in *rimb-1* mutants. This faster release in wild type thus must affect more efficient muscle activation, despite the larger currents in *rimb-1* mutants. To directly test this, we employed voltage imaging in muscle (**Fig. 4G**). The archaerhodopsin variant D95N shows voltage dependent fluorescence (Azimi Hashemi et al., 2019; Hochbaum et al., 2014; Kralj et al., 2012). Imaging Arch(D95N) before, during, and after 10 s photostimulation of cholinergic neurons revealed largely reduced muscle depolarization in *rimb-1* mutants, in line with the behavioral defects of *rimb-1* animals. Therefore, the summed outcome of fast synchronous SV fusions during evoked release is relevant for acute muscle depolarization, explaining the reduced muscle activation in *rimb-1* mutants, despite larger evoked currents. The latter could represent a homeostatic compensation in the mutant. However, we wondered if an alterntive explanation could be provided by the specific architecture of the *C. elegans* NMJ.

**Figure 4:**
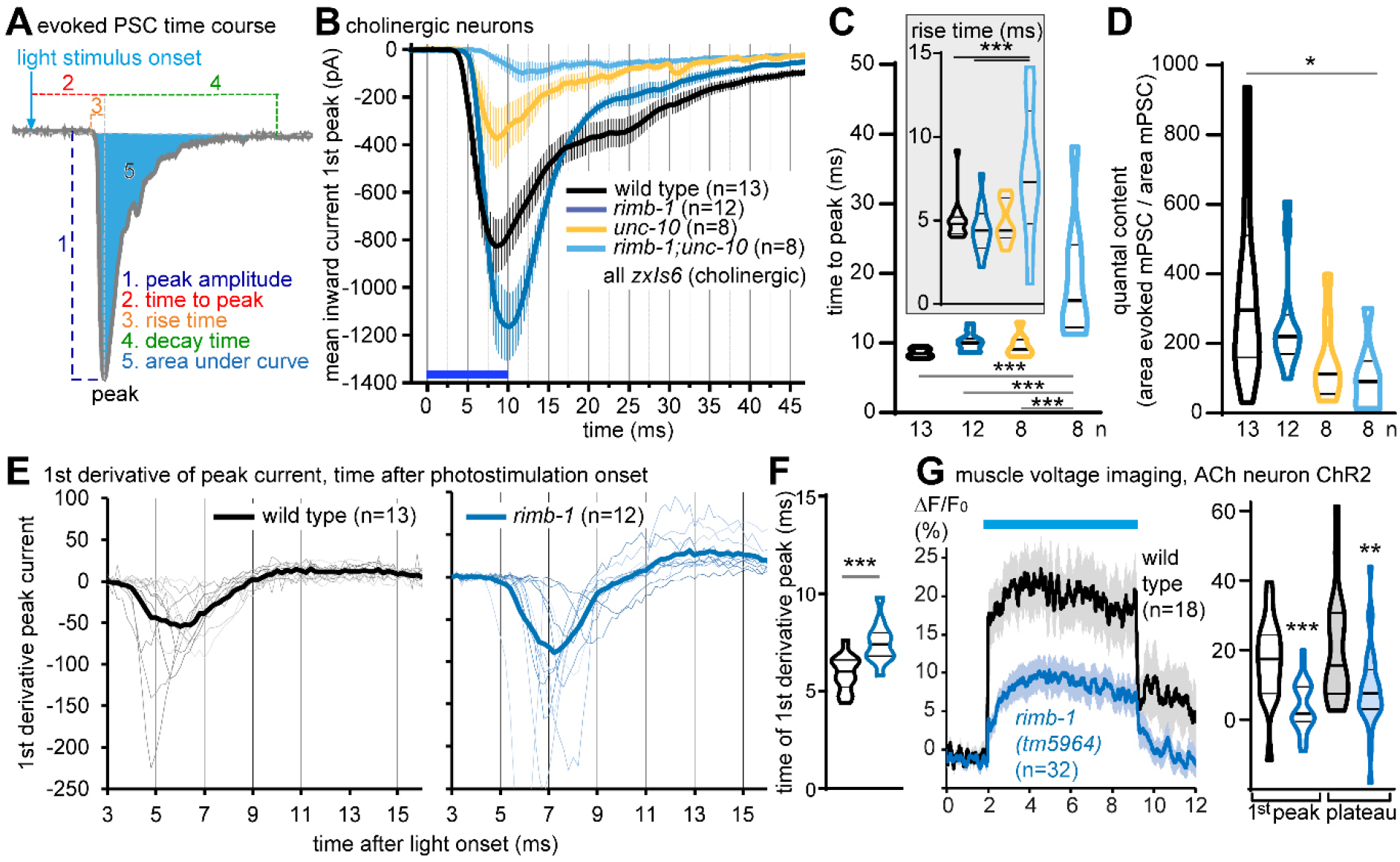
Parameters of evoked transmission reveal delayed postsynaptic current onset in *rimb-1* mutants. **A)** Analysis of ePSC properties, relative to the onset of the light stimulus (blue arrow). 1: peak amplitude; 2: time from light onset to peak; 3: rise time (from current onset to peak); 4: decay time (from peak to baseline); 5: area under the curve (charge transfer). **B)** High-resolution analysis of first light-evoked cholinergic peak current, mean ± s.e.m. **C)** Group data analysis of time to peak and rise time (inset). **D)** Quantal content, calculated as the area under the curve (ePSC) divided by the mPSC area under curve, from data in Fig. 3A. One-way ANOVA and Tukey test in C and Newman-Keuls test in D. **E)** Analysis of photo-evoked current slope (first derivative), to determine onset and peak current rise in individual (thin traces) and mean (thick line) stimulations, in wild type and *rimb-1* mutants. Time following photostimulus onset. **F)** Timing of the largest slope of the evoked currents, shown as median and 25/75 quartiles (thick and thin lines), min to max; unpaired t-test. **G)** Voltage imaging of muscle depolarization following cholinergic photostimulation. Muscles express the Arch(D95N) voltage sensor, fluorescence increases during photostimulation (blue bar); compared are wild type and *rimb-1* animals. Group data (peak response, plateau) shown as median and 25/75 quartiles (thick and thin lines), min to max. One-way ANOVA, Tukey test. Statistical significance given as *p<0.05; **p<0.01; ***p<0.001 in C, D, F, G.

### GABAergic ePSCs and summed NMJ transmission are reduced in *rimb-1* mutants

*C. elegans* body wall muscles are innervated not only by cholinergic, but also by GABAergic neurons (**Fig. 5A**), facilitating undulatory locomotion. Ventral (dorsal) cholinergic activity thus also evokes concomitant dorsal (ventral) GABAergic inhibition of muscle. Therefore, photostimulation of cholinergic neurons also evokes a GABAergic component. To address RIMB-1 function in GABAergic neurons specifically, we photostimulated them using ChR2 expressed from the *unc-47p* promoter (transgenes *zxIs3* or *zxIs9;* **Fig. 5B, C; S3A, B**). Over all time points, ePSCs were significantly smaller in *rimb-1; zxIs3* animals (**Fig. 5C, D**), while at the behavioral level, no significant difference to wild type was observed (**Fig. S3A, B**). Normalized GABA ePSCs did not reveal any difference over time (**Fig. S3C**). The time to peak was significantly delayed in *rimb-1* animals (**Fig. 5E, F**), while quantal content was unaltered (**Fig. 5G**). Thus, mutation of *rimb-1* delayed transmission in both cholinergic and GABAergic synapses, while it increased (ACh) or reduced (GABA) peak ePSCs. Therefore, cholinergic and GABAergic synapses may be differentially affected by *rimb-1* mutation. To distinguish this, we recorded spontaneous, as well as cholinergic neuron *(zxIs6*) evoked PSCs at a different holding potential (−10 mV), where GABA induced currents appear as outward (**Fig. S4A**). We observed a trend to reduced mPSC frequency for both ACh and GABA, and unaltered amplitudes (**Fig. S4B, C**). ePSCs were delayed, and smaller in *rimb-1* mutants (**Fig. S4D-H**).

**Figure 5:**
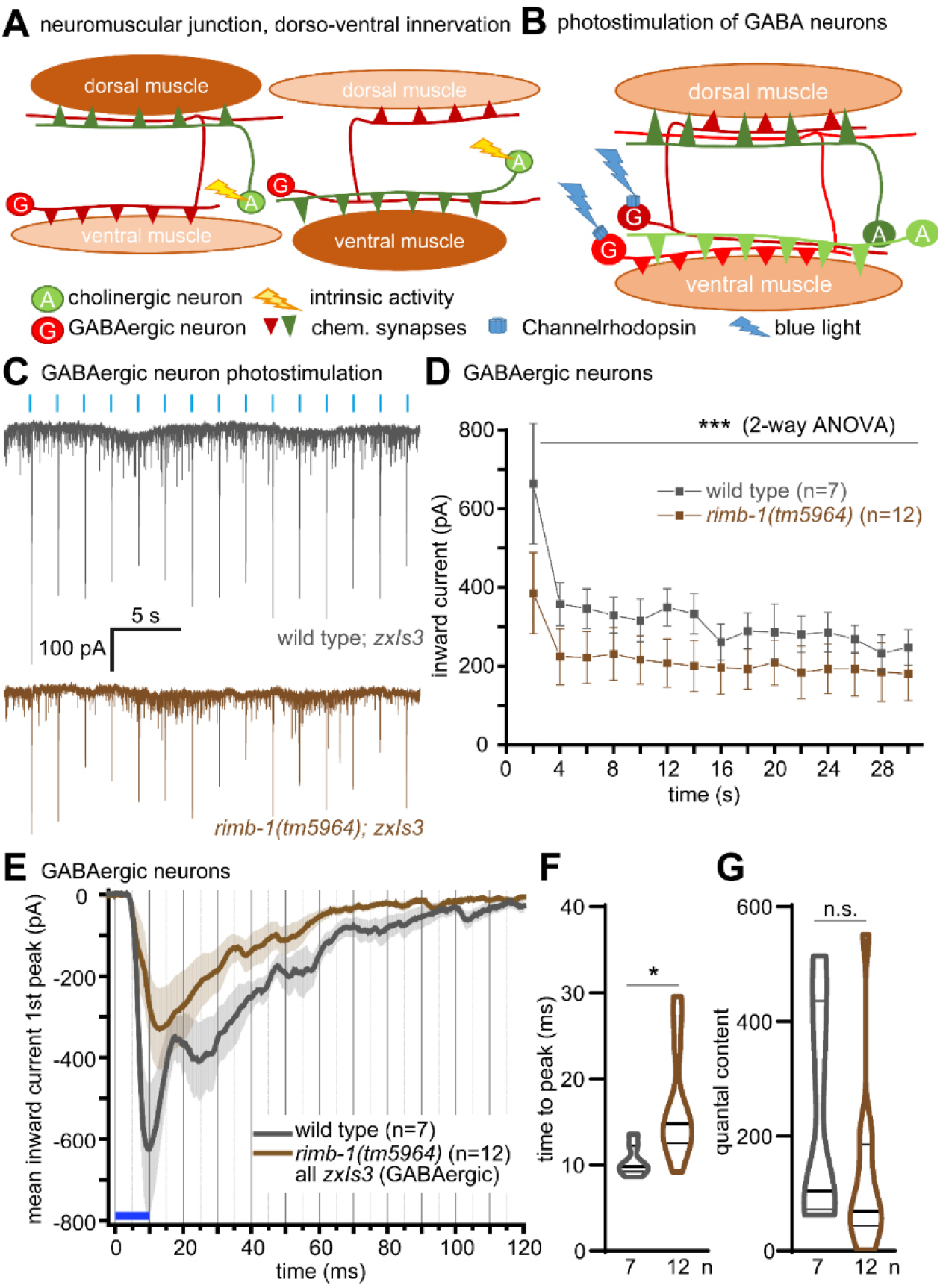
Lack of RIMB-1 reduces transmission from GABAergic motor neurons. **A)** Organization of the *C. elegans* NMJ: Reciprocal innervation of muscle by cholinergic (A, green) and GABAergic (G, red) neurons, and further contralateral innervation of GABAergic neurons by cholinergic neurons. Dorsal or ventral activity in cholinergic neurons (yellow flashes) causes concomitant GABAergic release and antagonistic inhibition, enabling undulatory locomotion. **B)** Specific optogenetic stimulation of GABAergic neurons (blue flashes). **C)** Animals carrying transgene *zxIs3,* expressing ChR2 in GABAergic neurons (promoter *unc-47p*) were photostimulated (0.5 Hz, 10 ms stimuli; blue tick marks), evoking postsynaptic currents; genotypes as indicated. Note that GABA induced currents appear as inward under the conditions used (−60 mV holding potential, specific pipette and bath solutions, see materials and methods). **D)** Repeated stimulation of GABAergic neurons (*zxIs3*), mean ± s.e.m. amplitudes of consecutive photostimuli. Two-way ANOVA. **E)** High-resolution analysis of first light-evoked GABAergic peak current, mean ± s.e.m. **F)** Group data analysis of time to peak, or quantal content **G)**, calculated as the area under the curve (ePSC) divided by the mPSC area under curve, from data in C. Shown as median and 25/75 quartiles (thick and thin lines), min to max. One-way ANOVA and Tukey test in F and Newman-Keuls test in G. Statistical significance given as *p<0.05; ***p<0.001.

This is more in line with a similar role of RIMB-1 in both cholinergic and GABAergic terminals. We wondered if differences in the two synapse types may be more apparent at higher stimulation regimes. Previously, it was shown that cholinergic synapses depress, while GABAergic synapse are facilitated at increased stimulation frequencies (Liewald et al., 2008; Liu et al., 2009). We therefore compared ePSCs during 2 Hz photostimulation in cholinergic *(zxIs6*) and GABAergic *(zxIs9*) neurons, in wild type and *rimb-1* mutants (**Fig. S5A-F**). At this stimulus frequency, both cholinergic and GABAergic synapses showed a fast run down during the first 3-4 pulses, and no obvious further depression. Importantly, in *rimb-1* mutants, cholinergic currents were slightly increased, while GABAergic currents were reduced (compared to wild type). Thus, the increased current in cholinergic synapses in *rimb-1* mutants does not originate from an increased activity in concomitantly activated GABAergic synapses.

### *rimb-1* defects are more pronounced in the absence of GABA transmission

To further probe the role of *rimb-1* in GABA transmission at the NMJ, we stimulated cholinergic neurons in *unc-47(e307*) mutants, lacking the vesicular GABA transporter (vGAT), and in *rimb-1; unc-47* double mutants. vGAT mutants contracted much more than wild type, since inhibitory transmission is abolished (**Fig. 6A**). In *rimb-1; unc-47* double mutants, the contractions were similar to wild type, i.e. in-between the single mutants. In addition, the double mutants showed a significant, progressive fatigue (i.e. SV recycling phenotype), as observed for *rimb-1* single mutants, though more pronounced. Thus, the lack of GABA transmission emphasizes the role of RIMB-1 in cholinergic transmission and SV recycling. mPSC frequency was significantly reduced in *unc-47* and (by about 80%) in *rimb-1; unc-47* double mutants (**Fig. 6B**), and likewise, mPSC amplitudes were significantly reduced in *rimb-1; unc-47* double mutants (**Fig. 6C**).

**Figure 6:**
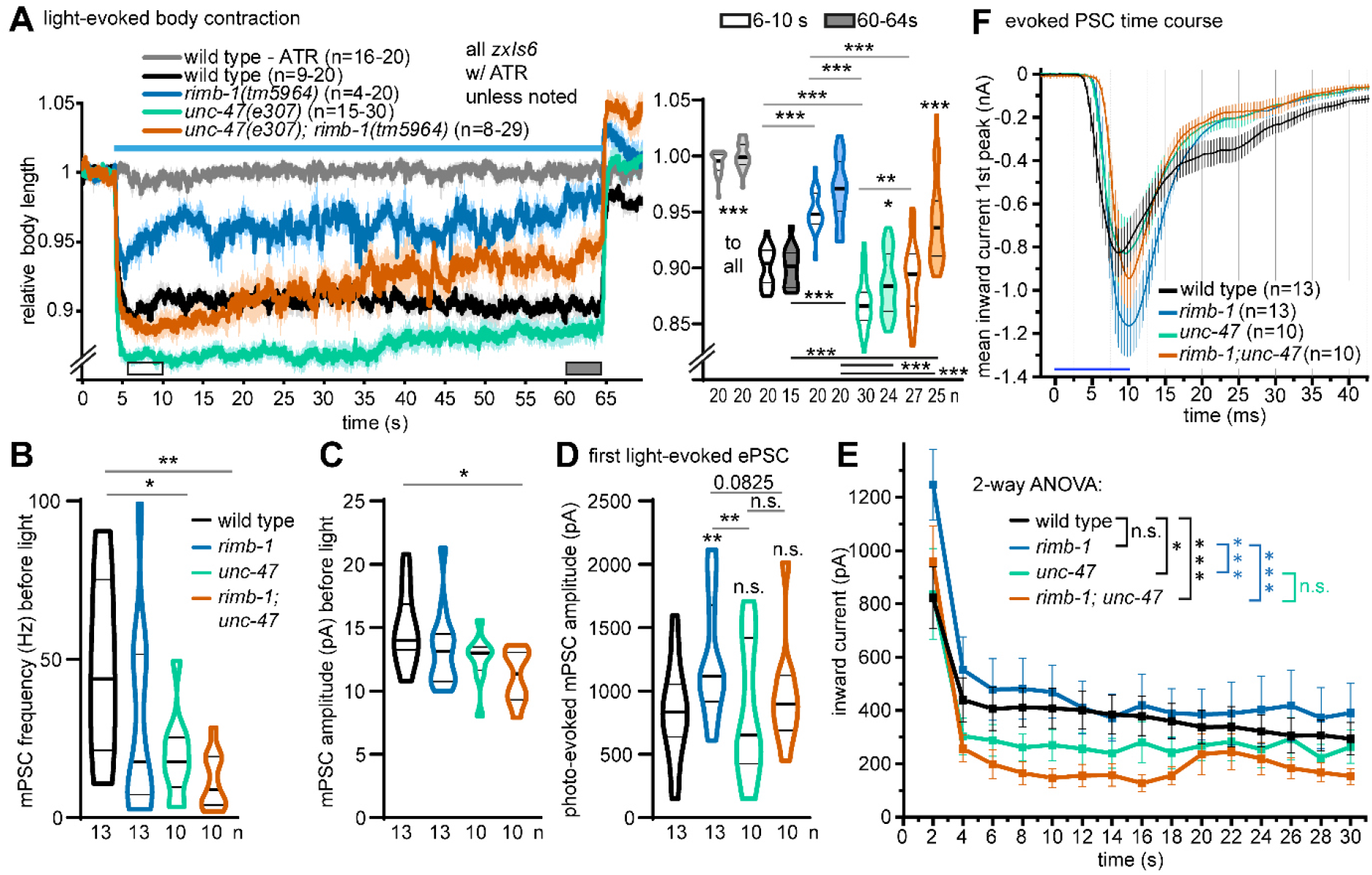
Lack of GABAergic transmission reverts increased cholinergic transmission in *rimb-1* mutants. **A)** Left panel: Body contraction after photostimulation of cholinergic neurons (*zxIs6*). Indicated genotypes and number of animals. Right panel: Group data for the 6-10 / 60-64 s time periods. Significant differences after one-way ANOVA with Tukey test (for same period) and two-way ANOVA with Bonferroni correction (for different time periods). **B)** mPSC frequencies and **C)** amplitudes during 30 s before first light stimulus for the indicated strains expressing ChR2 in cholinergic neurons (*zxIs6*). Data shown as median and 25/75 quartiles (thick and thin lines), min to max. n = number of animals. One-way ANOVA, Tukey test. **D)** Photoevoked PSC amplitude of first and **E)** consecutive photostimuli (mean ± s.e.m.). One-way ANOVA, Tukey test in D; two-way ANOVA with Fisher test in E. **F)** Time-course of ePSCs (first peak; blue bar: light stimulus); mean ± s.e.m. In A-E: Statistical significance given as *p<0.05; **p<0.01; ***p<0.001.

When we analyzed photoevoked cholinergic PSCs, we observed no difference for the comparison of wild type and *unc-47* mutants (**Fig. 6D-F**). This indicates that there is no relevant GABA contribution to muscle currents in these experiments, likely due to severing of cholinergic to GABAergic connections during the dissection. However, in the *rimb-1; unc-47* double mutant, the lack of GABA negatively affected the (increased) evoked amplitudes observed in *rimb-1* mutants (**Fig. 6D**): Amplitudes were not significantly different when compared to wild type. We suggest that the increased cholinergic ePSCs in *rimb-1* mutants reflect a homeostatic compensatory response to the temporally imprecise (delayed) cholinergic transmission (**Fig. 2, 4E-G**). In the absence of GABA, since the normal excitation-inhibition balance is disturbed (i.e., only excitation remains), the need for a homeostatic increase in cholinergic transmission in *rimb-1* animals is gone. Therefore, *rimb-1; unc-47* animals show normal evoked responses (**Fig. 6D**). In addition, *rimb-1; unc-47* mutants showed a strong rundown in 0.5 Hz stimulus trains, as in the contraction assay (**Fig. 6A, E, S3D**). Thus, synapses become more vulnerable to SV depletion in the absence of RIMB-1.

### RIMB-1 affects the location of SV fusion sites in the active zone

RIMB-1, together with UNC-10/RIM, aligns UNC-2/CaV2 VGCCs with primed SVs, ensuring that upon depolarization, Ca^2+^ microdomains are most effective in triggering SV fusion. Thus, the site of SV fusion, but possibly also the overall arrangement of the active zone (AZ), may be affected in *rimb-1* mutants. The *C. elegans* AZ harbors the dense projection (DP) at its center, where SVs are thought to be recruited for priming. We analyzed the ultrastructure of wild type and *rimb-1* mutant synapses by serial section transmission electron microscopy (TEM) (Kittelmann et al., 2013). 40 nm thin sections of high-pressure frozen (HPF), freeze-substituted and stained samples were analyzed, in resting or photostimulated animals (*zxIs6*) (Kittelmann et al., 2013; Steuer Costa et al., 2017; Yu et al., 2018). Following 30 s illumination in animals grown in presence or absence of all-*trans* retinal (ATR; thus rendering ChR2 active or inactive), total SVs, docked SVs, dense core vesicles (DCVs), and recycling-induced large vesicles (LVs) were quantified (**Fig. 7A-E**). Total SVs were slightly, but significantly reduced in *rimb-1* synapses, and photostimulation led to a more pronounced decrease in SV numbers in *rimb-1* mutants compared to wild type (**Fig. 7B**). This is surprising, since given the reduced fidelity of transmission in *rimb-1* mutants, one would expect reduced release and higher numbers of (docked) SVs. Possibly RIMB-1 is also required for SV docking. Indeed, there were significantly less docked SVs in *rimb-1* synapses, before and after photostimulation (**Fig. 7C**). DCV numbers were not affected by *rimb-1* or photostimulation (**Fig. 7D**; note DCV release in cholinergic neurons requires an increase in cAMP; Steuer Costa et al., 2017). Large vesicles, i.e. endosomes induced by photostimulation, were significantly increased in wild type photostimulated animals, but not in *rimb-1* synapses (**Fig. 7E**), in line with the reduced overall synaptic activity observed in behavioral experiments. DP size was not significantly different in wild type vs. *rimb-1* mutants, as it is in mammalian synapses (**Fig. S6A**; Acuna et al., 2016).

**Figure 7:**
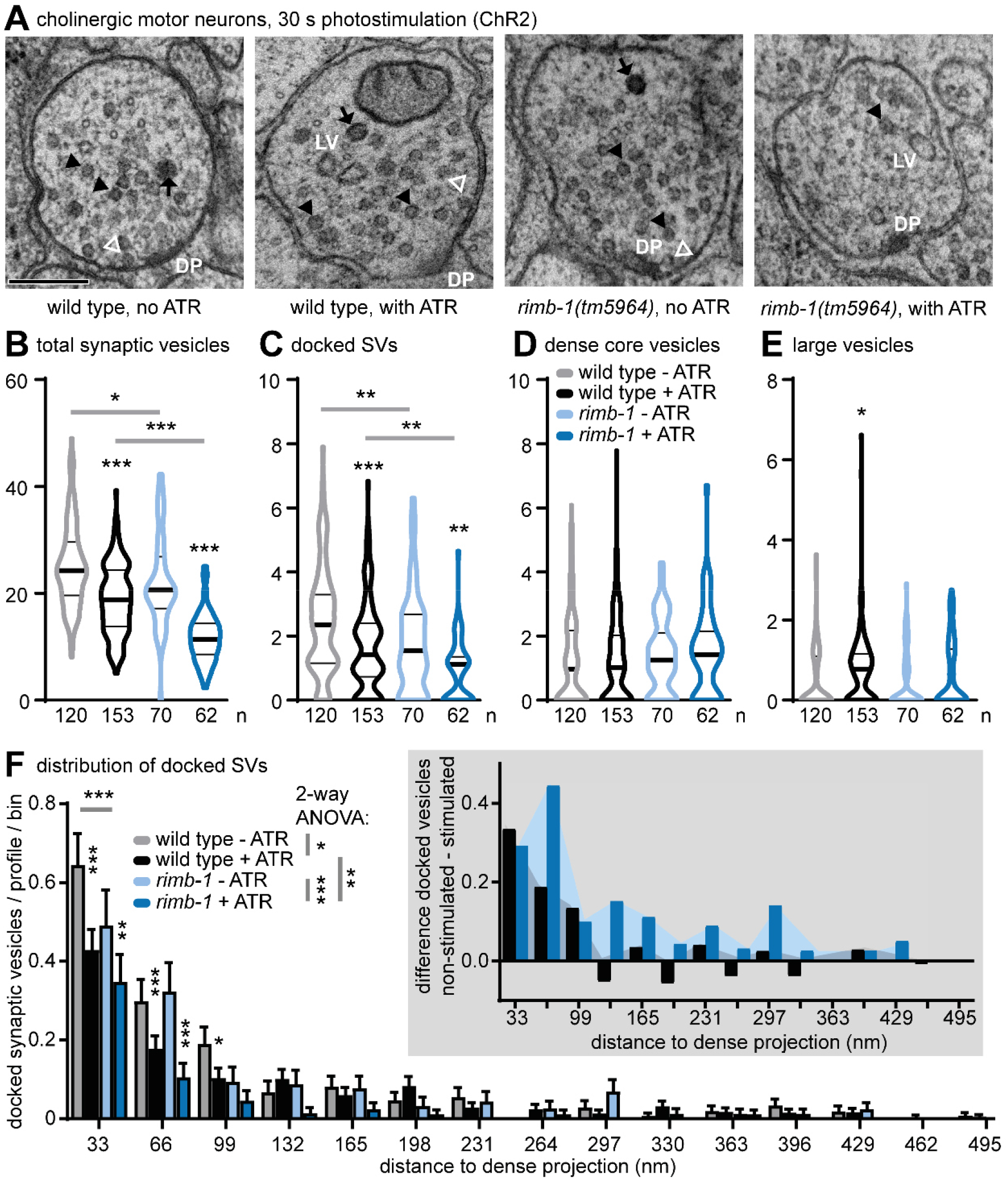
Electron microscopy analysis of photostimulated cholinergic *rimb-1* synapses. **A)** Transmission electron micrographs of cholinergic synapses expressing ChR2 (*zxIs6*), from animals that were high-pressure frozen following 30 s of photostimulation. 40 nm sections; genotype and ATR treatment indicated. Synaptic vesicles (SVs, closed arrowheads), docked SVs (open white arrowheads), dense core vesicles (DCVs, black arrows), large vesicles (LV, i.e. endosomes induced after photostimulation), and the dense projections (DP) are indicated. Scale bar is 200 nm. **B)** Quantification of all SVs, **C)** docked SVs, **D)** DCVs, and **E)** LVs, per profile. Data in B-E shown as median and 25/75 quartiles (thick and thin lines), min to max. One-way ANOVA with Kruskall-Wallis / Dunn’s multiple comparison test. **F)** The distribution of distances of docked SVs to the DP (along the plasma membrane) was analyzed in 33 nm bins, for the genotypes indicated. Two-way ANOVA / Tukey test (within bins, between groups). Inset: Difference in the number of docked SVs (normalized to 33 nm bin, no ATR) in each bin without photostimulation, minus with photostimulation (i.e. showing where SVs disappeared). See also **Fig. S7B**. In B-F: Statistical significance given as *p<0.05; **p<0.01; ***p<0.001.

Next, we analyzed the occurrence of docked SVs along the active zone membrane, i.e. their distribution relative to the DP (**Fig. 7F**). In non-stimulated synapses, docked SVs were most frequently found in close proximity to the DP, i.e. within 33 nm, and this leveled off to baseline between 132 and 165 nm distal to the DP. *rimb-1* mutants contained significantly fewer docked SVs in the 33 nm bin, but did not differ from wild type at other distances. This indicated that docked SVs distributed more distally in *rimb-1* mutants, possibly because SV tethering with the UNC-10/RIM scaffold is compromised (evident also from a cumulative distribution analysis; **Fig. S6B**). Upon photostimulation, wild type synapses fused SVs most effectively in close proximity to the DP, while in *rimb-1* mutants this occurred more distal to the DP (**Fig. 5F inset, Fig. S6B**). This is in line with putative mislocalization of UNC-2 VGCCs in *rimb-1* mutants, e.g. more distal to the DP, where SVs may experience a higher Ca^2+^ concentration than at the DP.

### Untethering P/Q type VGCCs affects transmission and counteracts the loss of RIMB-1

CaV2 VGCCs are tethered to docked SVs by RIM and RIMB-1 via multiple interactions, involving the SH3 and PDZ domains of RIMB-1 and RIM, respectively, which bind the UNC-2/CaV2 C-terminus (**Fig. 1B, 8A**).

**Figure 8:**
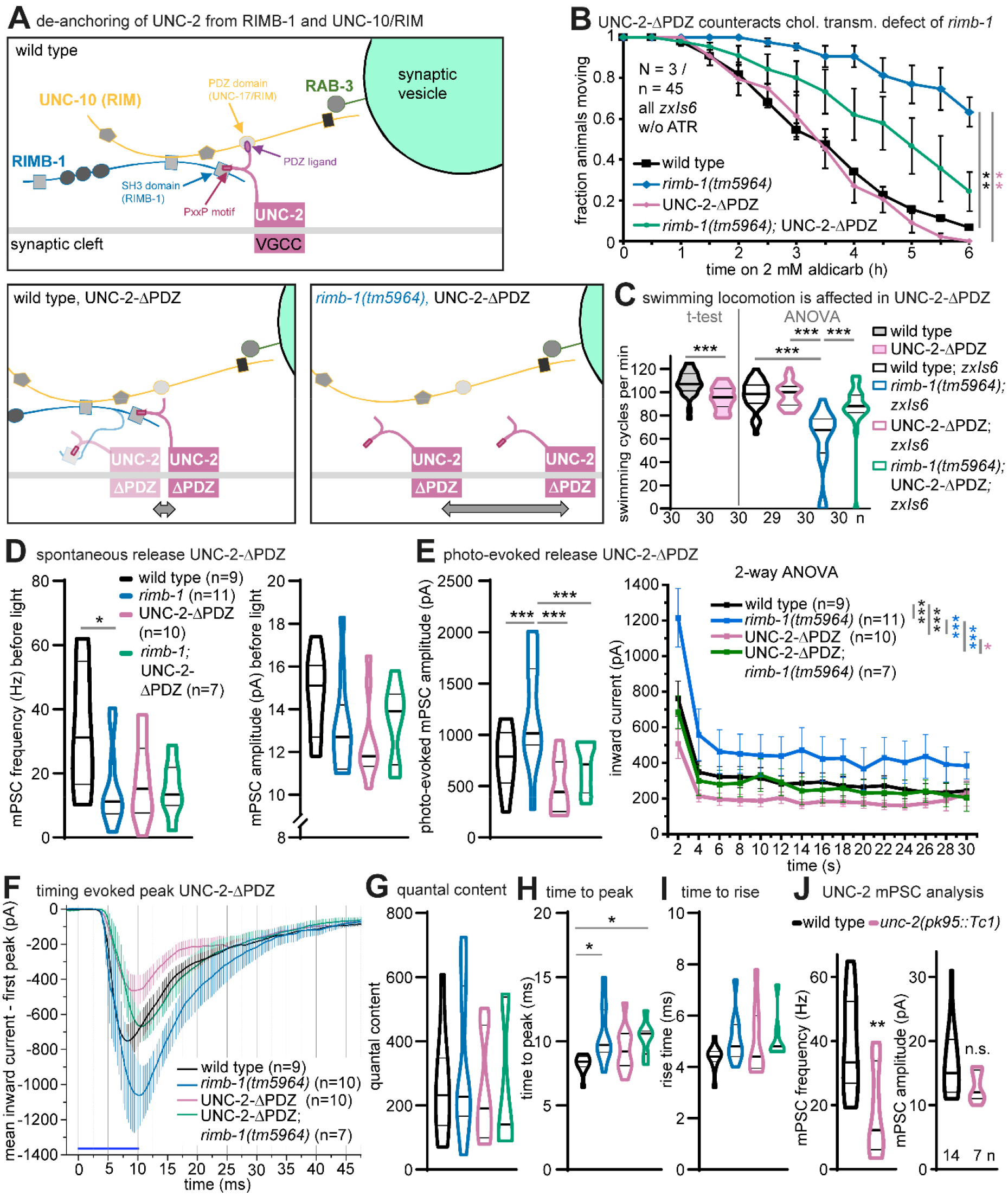
Untethering the CaV2 VGCC alters synaptic transmission, exacerbated by *rimb-1* deletion. **A)** Upper panel: Protein-protein interactions involving the CaV2(UNC-2) VGCC and RIMB-1. UNC-2 is tethered by interactions with RIMB-1 (PxxP motif bound by RIMB-1 SH3 domain) and RIM/UNC-10 (C-terminal PDZ ligand bound by UNC-10 PDZ domain). Lower left panel: Removing the PDZ ligand might loosen UNC-2 tethering at the docked SV site. Lower right panel: Removing RIMB-1 further untethers UNC-2-ΔPDZ. **B)** Aldicarb paralysis assay, mean ± s.e.m., 3 replicates, n=15 animals each, genotypes indicated. Log rank test, Bonferroni correction. **C)** Swimming assay, comparing locomotion rate in the indicated mutants, compared to wild type. Data shown as median and 25/75 quartiles (thick and thin lines), min to max. T-test or one-way ANOVA with Tukey test. n = number of animals tested. **D)** Electrophysiological analysis of UNC-2-ΔPDZ animals, compared to controls. mPSC frequency and amplitudes, before photostimulation. One-way ANOVA, Tukey test. **E)** Photoevoked (cholinergic neurons, transgene *zxIs6*) PSC amplitudes of first and consecutive stimuli. One-way ANOVA, Newman-Keuls test and two-way ANOVA, Fisher test. n = number of animals tested. **F)** Time-course of first ePSC (mean ± s.e.m.) and analysis of ePSC parameters **G-I)** quantal content, time to peak and time to rise. One-way ANOVA, Kruskall-Wallis test. **J)** Analysis of mPSC frequency and amplitudes in *unc-2* l.o.f. mutant; T-test. One-way ANOVA, Newman-Keuls test. n = number of animals. In B-E, G-J: Statistical significance given as *p<0.05; **p<0.01; ***p<0.001.

Thus far, we either analyzed the mild defect induced by the loss of RIMB-1 in the *tm5964* allele, or the very strong defect induced by loss of UNC-10/RIM. Phenotypes of the latter are strong, likely because they involve additional functions of RIM, not only UNC-2 tethering. We reasoned that untethering the UNC-2 channel may achieve an intermediate phenotype and exacerbate the *rimb-1* phenotype, when combined, allowing UNC-2 to diffuse even more (**Fig. 8A**). Thus the UNC-2 PDZ ligand (its C-terminal 10 amino acids) was removed by CRISPR-mediated genome editing; from now on we refer to this mutant as UNC-2-ΔPDZ. First, we analyzed cholinergic transmission in UNC-2-ΔPDZ animals and in *rimb-1;* UNC-2-ΔPDZ double mutants (**Fig. 8B**). UNC-2-ΔPDZ *per se* had no phenotype in the aldicarb assay, however, it appeared to counteract rather than exacerbate the resistance phenotype of *rimb-1* mutants. UNC-2-ΔPDZ animals were slightly, but significantly affected for swimming. Again, the *rimb-1;* UNC-2-ΔPDZ double mutants were somewhat rescued when compared to the *rimb-1* mutants, and this was similarly found for ChR2-evoked body contractions (**Fig. S7A**). Next, we analyzed synaptic transmission by electrophysiology in UNC-2-ΔPDZ and *rimb-1;* UNC-2-ΔPDZ animals, before and during photostimulation of cholinergic neurons (**Fig. 8D-I**). UNC-2-ΔPDZ did not significantly reduce spontaneous release (mPSC rate), but it reverted the significantly reduced mPSC rate of *rimb-1* animals (**Fig. 8D**), while mPSC amplitudes were not altered. Thus, untethering of UNC-2 is not strongly deleterious for spontaneous activity, and it may allow the VGCC to trigger SVs that it cannot efficiently reach in the *rimb-1* mutant, when it is still tethered to UNC-10/RIM. However, upon photostimulation, UNC-2-ΔPDZ animals exhibited strongly reduced ePSC amplitudes when compared to *rimb-1* mutants and to wild type (**Fig. 8E**). This indicates that UNC-2-ΔPDZ supports spontaneous release, but fails to do so during evoked activity. Possibly, the strong optogenetic stimulus requires tight coupling of UNC-2 and docked SVs. UNC-2-ΔPDZ, and particularly *rimb-1;* UNC-2-ΔPDZ double mutants, were delayed in their time to peak (**Fig. 8F, H**), in line with an increased distance of the VGCC to the docked SV, while the rise time was unaffected (**Fig. 8I**). Quantal content was unaltered in this set of mutants (**Fig. 8G**).

We compared effects of untethering vs. absence of UNC-2. In *unc-2(pk95::Tc1*) loss-of-function mutants, mPSC frequency was largely reduced, but mPSC amplitude was unaltered (**Fig. 8J**), similar to the UNC-2-ΔPDZ mutant and the *rimb-1;* UNC-2-ΔPDZ double mutants (**Fig. 8D; S7B**). In contraction assays, *unc-2(pk95::Tc1*) mutants showed largely reduced effects and were most similar to *rimb-1(tm5964*) mutants (**Fig. S7A, C**). Thus, the absence of the channel is essentially comparable to the loss of its tethering.

### Acute degradation of the CaV2 channel β-subunit CCB-1 reduces synaptic transmission

Because the *rimb-1* and *unc-2* mutations may induce long-lasting compensatory effects, we wanted to acutely eliminate UNC-2 function, and to compare this to the completely untethered UNC-2-ΔPDZ in *rimb-1* background. To this end, we used a photosensitive degron (PSD), which can effectively degrade synaptic proteins within 45-60 min of mild blue illumination (Hermann et al., 2015). The PSD was fused to the C-terminus of the essential VGCC β-subunit CCB-1 by CRISPR-editing (**Fig. 9A**). CCB-1 is required for VGCC biogenesis and function (Laine et al., 2011). Photodegradation is expected to disrupt the CCB-1 protein and to either inactivate the VGCC complex, or to destabilize it and cause its disruption. 1 h photodegradation of CCB-1::PSD caused aldicarb resistance, though only in the absence of GABAergic signaling in *unc-47* mutants (likely because of an excitation-inhibition imbalance in the CCB-1::PSD strain; **Fig. 9B**). Furthermore, it significantly reduced swimming, compared to illuminated wild type animals, and this effect recovered after 24h (**Fig. 9C**). The mPSC frequency was significantly decreased by about 75% in CCB-1::PSD animals after 1 h blue light illumination, confirming acute degradation of CCB-1 (**Fig. 9D**), and was comparable to the UNC-2 deletion (**Fig. 8J, S7B**). Likewise, it did not significantly affect mPSC amplitude. Thus, UNC-2 function is affected by the acute degradation of the CCB-1 β-subunit, either through an effect on channel properties, or by affecting stability of the channel in the plasma membrane, or both. In sum, untethering of CaV2/UNC-2 could partially balance the absence of RIMB-1. Possibly, the complete untethering allows diffusion of the CaV2 channel near docked SVs that are in the periphery of the AZ. Since the deletion of UNC-2/CaV2 does not eliminate all transmission, this implies that other VGCCs partially compensate for function at the NMJ, most likely EGL-19/CaV1 L-type VGCCs, as was recently shown (Liu et al., 2018; Tong et al., 2017).

**Figure 9:**
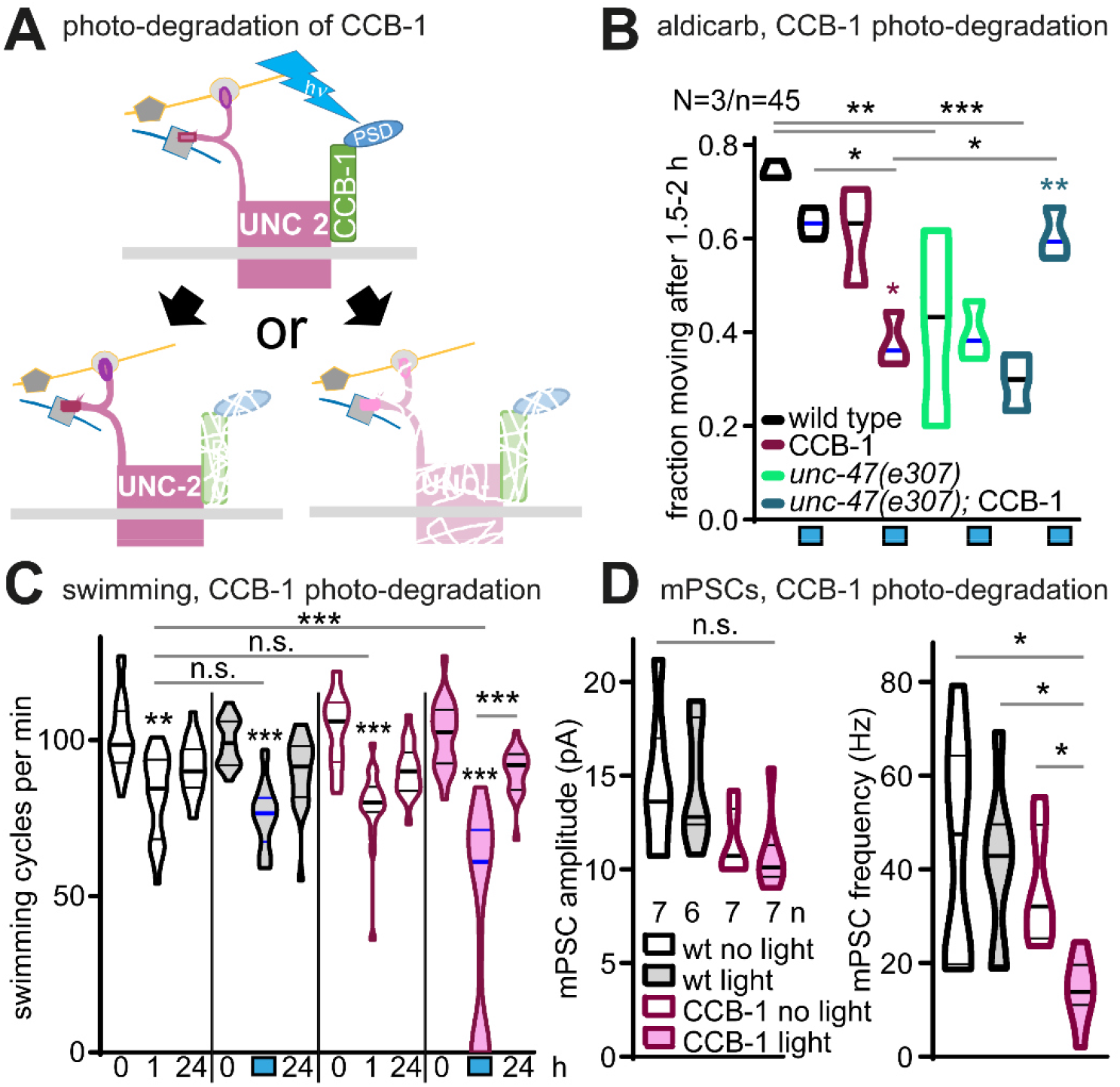
Acute photodegradation of the CaV2 β-subunit CCB-1 affects behavior and reduces synaptic transmission. **A)** Acutely inactivating (or destabilizing) UNC-2 by photodegradation of the β-subunit, CCB-1, tagged with a C-terminal photo-sensitive degron (PSD). **B)** Animals (genotype and/or treatment as indicated: CCB-1: PSD-tagged version; *unc-47* vGAT mutants; blue boxes: photostimulation for 1h) were analyzed in aldicarb (1 mM) paralysis assay. Data averaged from the 1.5-2 h time points. Two-way ANOVA with Tukey test. **C)** Swimming assay in CCB-1::PSD animals before and after photodegradation (blue bar, shaded data sets) and following 24 h recovery. Compared to wild type controls, and non-illuminated animals (non-shaded data sets). One way ANOVA with Bonferroni test within data sets (light and no light) and two-way ANOVA to compare datasets without and with illumination. **D)** mPSC analysis, frequency and amplitude, without and with CCB-1::PSD photo-degradation (1h), and wild type controls.

### UNC-2/CaV2 synaptic density is reduced in the absence of RIMB-1 or the β-subunit

The de-anchoring of the UNC-2 channel via elimination of the RIMB-1 protein and the PDZ ligand at the UNC-2 C-terminus affect (i.e. increase) its mobility in the synapse, and thus likely diminish its location in vicinity of docked SVs (i.e. move it away, on average). Yet, it may also affect its overall amount in the active zone membrane. To address this, we imaged GFP::UNC-2 in nerve ring synapses (**Fig. 10A**; GFP was inserted by CRISPR in the N-terminal domain of UNC-2; strain kindly provided by Mei Zhen; Gao et al., 2018). Further, the UNC-2 PDZ ligand was deleted in this strain, and we compared both strains in wild type and *rimb-1(tm5964*) background, assessing overall nerve ring fluorescence level as well as synaptic puncta intensity and size (**Fig. 10B**). Fluorescence levels were significantly reduced in animals lacking both RIMB-1 and the UNC-2 PDZ ligand, compared to all other genotypes. This could be due to less clustering, or to overall reduction of the (synaptic) expression level of the CaV2/UNC-2. The fluorescence intensity of UNC-2 clusters was also significantly reduced without RIMB-1, and even more so by the additional deletion of the UNC-2 PDZ ligand. Cluster size was not significantly different. Thus, loss of RIMB-1 and UNC-2 anchoring reduces synaptic levels of the channel, explaining the reduced synaptic function. Alternatively, UNC-2 trafficking to the synapse is inhibited, or its localization is more diffuse.

**Figure 10:**
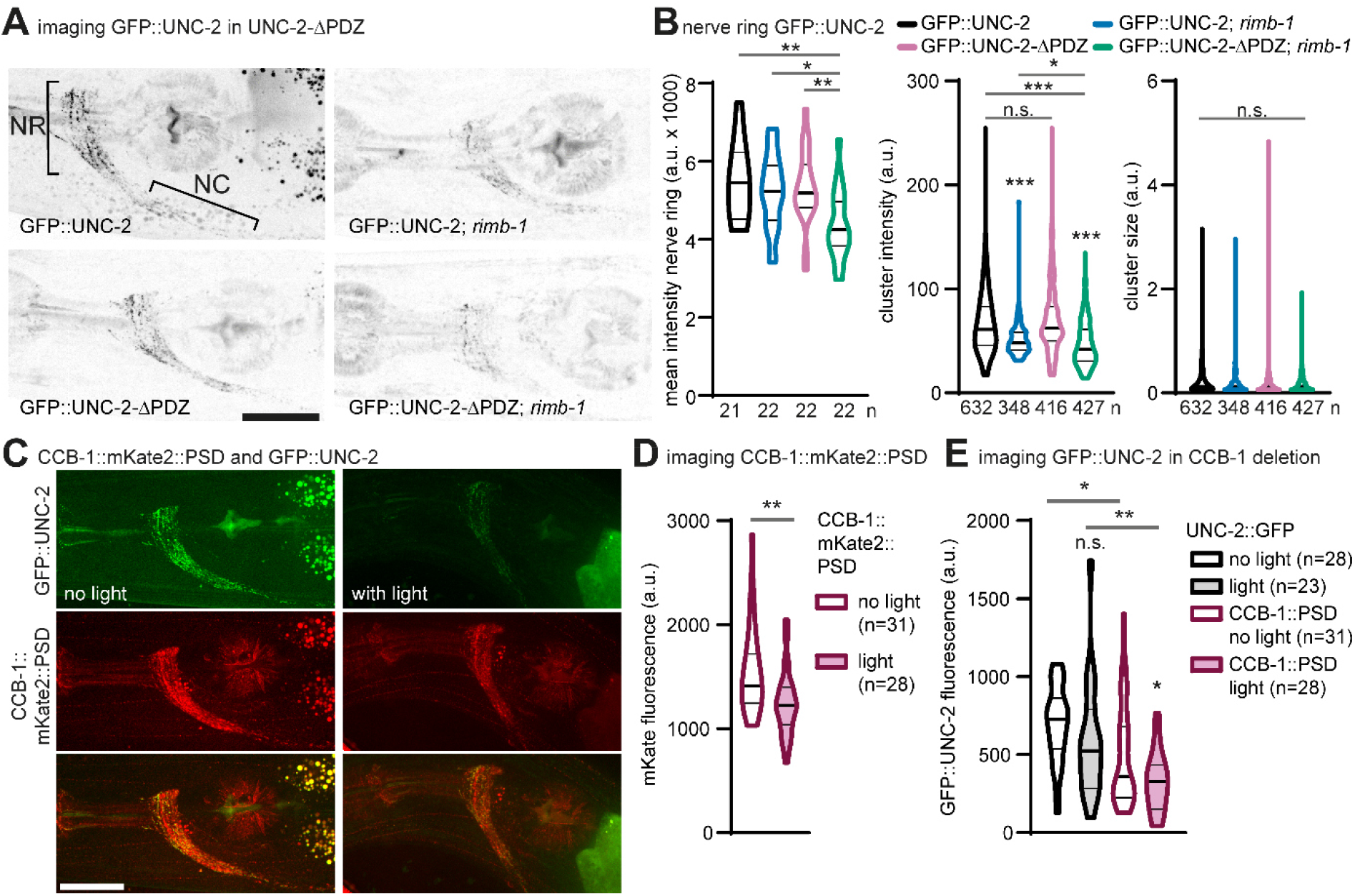
CaV2/UNC-2 channels are reduced at synapses in *rimb-1* mutants and after acute photodegradation of the CCB-1 β-subunit. **A)** Confocal z-projections of fluorescence of GFP::UNC-2 in the nerve ring (NR) and anterior ventral nerve cord (NC). Anterior is to the left, scale bar = 20 μm, genotypes as indicated. **B)** Quantification of fluorescent puncta (synaptic UNC-2 clusters) as shown in A, mean intensity of nerve ring (n= number of animals), mean cluster fluorescence and size (n= number of puncta). Data shown as median and 25/75 quartiles (thick and thin lines), min to max. One-way ANOVA with Tukey test or Kruskal-Wallis / Dunn’s test. **C)** Confocal stacks of nerve ring fluorescence of GFP::UNC-2 (green) and CCB-1::mKate2::PSD (red), and overlay, before and after blue light activation of the PSD for 1 h; scale bar = 20 μm. **D)** CCB-1::mKate2::PSD fluorescence in the nerve ring, before and after blue light stimulation; n = number of animals analyzed. **E)** Quantification of GFP::UNC-2 in controls and in animals expressing CCB-1::mKate2::PSD, before and after 1 h blue illumination. Data shown as median and 25/75 quartiles (thick and thin lines), min to max. T-test or one-way ANOVA with Newman-Keuls test. In B, D: Statistical significance given as *p<0.05; **p<0.01; ***p<0.001.

We also assessed the effects of acute photodegradation of the CCB-1::PSD β-subunit on UNC-2 abundance. GFP::UNC-2 and CCB-1::mKate2::PSD co-localized in synaptic puncta, although CCB-1 was more abundant and present also at more peripheral synaptic regions (**Fig. 10C**). Fluorescence of CCB-1::mKate2::PSD was significantly reduced after 1 h of blue illumination (**Fig. 10D**), while GFP::UNC-2 signal was not affected by the same blue light treatment (**Fig. 10E**). However, combination with CCB-1::PSD reduced GFP::UNC-2 levels in the dark, and, even more, in response to blue light. Thus, absence of the β-subunit affected UNC-2 function and synaptic density in a similar way as the absence of RIMB-1 and the UNC-2 PDZ ligand, most likely affecting the close proximity of the P/Q-type VGCC to docked SVs.

## Discussion

The fidelity and robustness of synaptic transmission depends on the precise morphology and spatial organization of the active zone (Sudhof, 2013). Thus, mutations in proteins that orchestrate the anchoring and tight apposition of docked SVs and CaV2 VGCCs are likely to negatively affect this process. Also, such proteins are probable targets for regulation of synaptic transmission, and could be utilized to set the excitation-inhibition balance also in complex neural circuits (Huang et al., 2019; Liu et al., 2018). Here, we characterized in detail the function of the *C. elegans* RIMB-1 protein, as well as its (functional) interaction with the CaV2/UNC-2 VGCC, and determinants of VGCC localization at NMJ synapses, in cholinergic and GABAergic neurons.

The lack of RIMB-1 function in *C. elegans* synapses had a mild phenotype on behavior (swimming) and caused some resistance to the ACh esterase inhibitor aldicarb, demonstrating cholinergic deficits. In response to optogenetic stimulation, the defect was also observable, however, lack of UNC-10/RIM had much more severe defects. The phenotypes of the double mutant were dominated by the *unc-10* defects, somewhat contradictory to an earlier report (Kushibiki et al., 2019), that assessed UNC-2::GFP overexpressed from a transgene. The situation in worms differs from that of the *Drosophila* NMJ, where *drbp* mutants had more severe phenotypes than *rim* mutants, depending on whether baseline or plasticity inducing conditions were studied (Liu et al., 2011; Muller et al., 2015).

Electrophysiology surprisingly showed increased transmission in *rimb-1* mutants in response to cholinergic stimulation. However, voltage imaging demonstrated reduced depolarization in *rimb-1* animals, in line with the behavioral defects. The likely reason for the decreased depolarization in response to cholinergic neuron activation is the reduced acuteness of release, which is delayed and likely not as synchronous as required to induce muscular action potentials (Liu et al., 2013). Therefore, the neurons try to balance this by increased output of ACh, leading to increased postsynaptic current. However, as this occurs at the wrong time, it can only partially compensate the *rimb-1* mutant deficit. In support of this hypothesis, we found that in mutants lacking GABA, the *rimb-1* mutant neurons performed much worse. Likely, this is because the lack of inhibition leads to a lower homeostatic drive, thus *rimb-1* cholinergic neurons do not upregulate their output as much. Overall, our findings show a differential importance of RIMB-1 in cholinergic vs. GABAergic synapses. While cholinergic synapses compensate the lack of RIMB-1 by increasing the ACh output, GABAergic synapses are weakened without any obvious homeostatic response. Such differential functions are also found in different central synapses in the mouse (Brockmann et al., 2019). In *C. elegans*, a gain-of-function mutation in UNC-2/CaV2 can induce increased cholinergic, but reduced GABA transmission, through effects on GABA_A_ receptors, or rather, GABA synapse numbers (Huang et al., 2019). Here, we probed post-synaptic nAChRs through levamisole paralysis assays, which did not show a defect in *rimb-1* mutants. We probed GABAergic synapses by direct ChR2-induced stimulation, which showed reduced currents in *rimb-1* mutants. Spontaneous GABA transmission showed reduced frequency but unaltered amplitudes in *rimb-1* animals. An additional layer of complexity at the *C. elegans* NMJ is given by the feedback inhibition that GABAergic neurons can elicit to cholinergic motor neurons via the AVA premotor interneurons (Liu et al., 2020). Also this way, reduced synaptic transmission due to the lack of RIMB-1 may affect the summed outcome of NMJ transmission.

Mislocalization of CaV2/UNC-2 leads to lower ACh release as fewer SVs are accessed by Ca^2+^. This is supported by our analysis of spontaneous release (mPSCs), and in behavioral assays. However, if CaV2/UNC-2 channels can rather freely diffuse in the synaptic membrane, they may also affect SVs that are docked in regions where they would normally not fuse. When optogenetic stimulation is used, which is much stronger than physiological activity during spontaneous release (Liewald et al., 2008; Liu et al., 2009), the larger Ca^2+^ domains could reach more docked SVs, distal to the active zone, where fusion events can still be observed (Watanabe et al., 2013). This slows down transmission, as we show, but may at the same time increase the amount of ACh released. The time from an optogenetic stimulus to reaching ePSC peak was delayed in both cholinergic and GABAergic *rimb-1* mutant synapses; thus, as in other systems (Acuna et al., 2016), RIMB-1 function is required for faithful and fast transmission. Quantal content was not reduced in *rimb-1* mutants, thus even if fewer SVs were released in a fast manner, overall the synapse was able to fuse normal numbers of SVs, only over a prolonged time. This indicates that Ca^2+^ levels in the vicinity of SVs rise more slowly in *rimb-1* mutants, but eventually reach sufficient levels to fuse SVs. As our ultrastructural analysis showed, the site of release was more remote from the dense projection, likely because CaV2/UNC-2 channels moved to more distal positions relative to the DP in the active zone membrane.

The deletion of the C-terminal PDZ ligand of the UNC-2 channel, by which it is anchored to RIM, had in part similar effects as the *rimb-1* mutation (mPSC frequency), in part it was less (behavior), or more severe (ePSC amplitude, evoked body contraction). In combination, the phenotypes of *rimb-1* and UNC-2-ΔPDZ were exacerbated or counteracted each other. This is difficult to reconcile, but may again be explained by the different degree of UNC-2/CaV2 de-anchoring, complex signaling in the motor circuit, spontaneous versus evoked activity and the need for precise VGCC localization for the latter. The photodegradation of the CCB-1::PSD β-subunit, which strongly reduced mPSC frequency and thus likely induced an acute functional elimination of the CaV2 channel complex, had a similar effect as its de-anchoring. Both manipulations led to a similar reduction in GFP::UNC-2 signal at nerve ring synapses, which may be due to destabilization and partial loss of the channel; yet also a re-localization away from synaptic regions into the axonal plasma membrane appears possible. To rigorously distinguish these possibilities, further work is required. In a recent publication, RIMB-1 function was assessed with respect to its role in building the synaptic cytomatrix/DP and in anchoring CaV2 channels (Oh et al., 2021). These authors found a contribution of RIMB-1 and a larger effect of UNC-10/RIM mutations on CaV2/UNC-2 abundance at synapses.

The effect of eliminating all CaV2 channels on synapse structure was analyzed in mammalian synapses (Held et al., 2020). Overall synapse structure was not affected, although the CaV2 β-subunit lost its localization to synaptic regions. This is in line with our findings, where loss of CCB-1 also affected UNC-2 synaptic location and/or abundance, thus these proteins may mutually influence each other. The authors also analyzed the deletion of the entire C-terminus of murine CaV2.1, which demonstrated that it is required for channel targeting to the synapse, and for triggering of efficient SV release. However, in mice, removing only the C-terminal PDZ ligand had no functional effects on its own, apart from a slightly lowered CaV2 abundance, but had to be combined with removal of the CaV2 RIM-BP binding site to evoke effects. This latter situation resembles our combination of *rimb-1* and UNC-2-ΔPDZ. Another study of the CaV2.1 C-terminus, however, does not observe alterations in VGCC function in mice when the RIM and RIM-BP binding sites are simultaneously removed (Lubbert et al., 2017). Thus, there may be species-specific differences in the regulation of the CaV2 channel and its synaptic localization by binding partners.

Overall our work confirms the general evolutionary conservation of RIM-BP function in *C. elegans*, and provides new information on how the protein regulates synaptic transmission through accurate placing of CaV2 channels in the vicinity of docked SVs. It further provides information on how loss of this protein affects excitation-inhibition balance in complex circuits, implying that regulation of this balance may be achieved *via* RIMB-1 by differentially affecting each synapse type, in this case cholinergic and GABAergic motor neurons. In the future, cell-type specific determinants of RIMB-1 function, e.g. through posttranslational modification or changes in expression level, will have to be revealed. Last, our work shows that de-anchoring the CaV2 channel from the active zone dense projection has effects similar to its elimination. The remaining transmission must be mediated by other VGCCs, most likely CaV1/EGL-19.

## Materials and Methods

### Strains

*C. elegans* wild type Bristol N2 and mutant strains were maintained on NGM plate seeded with *E. coli* OP50 as described earlier (Brenner, 1974). Young adult hermaphrodites cultured at 20°C were used. Mutant alleles and integrated transgenes used in this study:

*rimb-1(tm5964*)
**CB307**: *unc-47(e307*)
**KG4995:** *rimb-1 (ce828*)
**NL735**: *unc-2(pk95::Tc1)X*
**NM1657**: *unc-10(md1117*)
**ZM9583**: *unc-2(hp858[GFP::UNC-2]*) (GFP knock-in in genomic location of *unc-2*)
**ZX426**: *zxIs3[punc-47::ChR2(H134R)::yfp; lin-15+]*
**ZX460**: *zxIs6[punc-17::ChR2(H134R)::yfp; lin-15+]V*
**ZX531**: *unc-47(e307)III; zxIs6[punc-17::ChR2(H134R)::yfp; lin-15+]*
**ZX668:** *zxIs9[punc-47::ChR2(H134R)::YFP; unc-119+]*
**ZX1605**: *unc-2(pk95::Tc1*); *zxIs6[punc-17::ChR2(H134R)::yfp; lin-15+]*
**ZX1954**: *zxIs120[pmyo-3::Arch(D95N)::2xmyc; pmyo-3::CFP]; zxIs6[punc-17::ChR2(H134R)::yfp; lin-15+]*
**ZX2025**: *rimb-1(tm5964)III; zxIs6[punc-17::ChR2(H134R)::yfp; lin-15+]*
**ZX2271**: *ccb-1(zx3[ccb-1::mKate2::PSD]*) (CRISPR knock-in of mKate2 with a photosensitive degron (PSD))
**ZX2320**: *ccb-1(zx3[ccb-1::mKate2::PSD]); zxIs5[punc-17::ChR2(H134R)::yfp; lin-15+]*
**ZX2321**: *ccb-1(zx3[ccb-1::mKate2::PSD]); unc-2(hp858[GFP::UNC-2]*)
**ZX2340**: *unc-10(md1117); rimb-1(tm5964*)
**ZX2412**: *unc-10(md1117); rimb-1(tm5964); zxIs6[punc-17::ChR2(H134R)::yfp; lin-15+]*
**ZX2413**: *unc-10(md1117); zxIs6[punc-17::ChR2(H134R)::yfp; lin-15+]*
**ZX2474**: *zxEx1137[primb-1::GFP; punc-17::mCherry]*
**ZX2574**: *rimb-1(tm5964)III; zxIs3[punc-47::ChR2(H134R)::yfp; lin-15+]*
**ZX2562**: *rimb-1(tm5964)III; oxIs353[pmyo-3::ChR2::mCherry::unc-54 3’-UTR; lin15+]V*
**ZX2573**: *ccb-1(zx3[ccb-1::mKate2::PSD]); unc-47(e307) III.*
**ZX2584**: *rimb-1(tm5964); zxIs120[pmyo-3::Arch(D95N)::2xmyc; pmyo-3::CFP]; zxIs6[punc-17::ChR2(H134R)::yfp; lin-15+]*
**ZX2623/PHX1749**: *unc-2(syb1749[UNC-2-ΔPDZ]*) (UNC-2-ΔPDZ CRISPR deletion of 10 C-terminal amino acids)
**ZX2624**: *unc-2(syb1749[UNC-2-ΔPDZ]); zxIs6[punc-17::ChR2(H134R)::yfp; lin-15+]*
**ZX2625**: *unc-2(syb1749[UNC-2-ΔPDZ]); rimb-1(tm5964)III; zxIs6[punc-ZX2626: rimb-1(tm5964)III; unc-47(e307)III; zxIs6[punc-17::ChR2(H134R)::yfp; lin-15+]*
**ZX2627**: *snb-1(md247)V; oxIs353[pmyo-3::ChR2::mCherry::unc-54 3’-UTR; lin15+]V*
**ZX2691**: *unc-2(syb2088[UNC-2-ΔPDZ]); unc-2(hp858[GFP::UNC-2]*)
**ZX2757**: *unc-2(syb1749[UNC-2-ΔPDZ]*) 3x outcrossed
**ZX2758**: *unc-2(syb1749[UNC-2-ΔPDZ]); zxIs6[punc-17::ChR2(H134R)::yfp; lin-15+]*
**ZX2759**: *unc-2(syb1749[UNC-2-ΔPDZ]); rimb-1(tm5964)III; zxIs6[punc-17::ChR2(H134R)::yfp; lin-15+]*
**ZX2691/PHX2088**: *unc-2(syb2088[UNC-2-ΔPDZ]); unc-2(hp858[GFP::UNC-2]*) (UNC-2-ΔPDZ CRISPR deletion of 10 C-terminal amino acids in GFP::UNC-2 insertion)
**ZX2824**: *rimb-1(tm5964) III; unc-2(syb2088[UNC-2-ΔPDZ]); unc-2(hp858[GFP::UNC-2]*)
**ZX2825**: *rimb-1(tm5964) III; unc-2(hp858[GFP::UNC-2]*)
**ZX3112:** *rimb-1(tm5964); zxIs9[punc-47::ChR2(H134R)::YFP; unc-119+]*

### Molecular biology

To investigate the expression pattern of RIMB-1, we created *a rimb-1p::GFP* plasmid. The *rimb-1p* promoter (3 kb upstream of ATG) was amplified by PCR from genomic DNA using primers oBJ76 (TAGCTCTTCCAGCGAGAGGACCTCCTCCTC) and oBJ77 (AGCGCTGTTGGTGACTAGGTGGTCC), was inserted into vector pCS179 *(flp-1p(trc)::GFP*) containing GFP, described in (Oranth et al., 2018).

The strain ZX2271 *(ccb-1(zx3[ccb-1::mKate2::PSD])*) was generated using a CRISPR knock-in strategy (Dickinson et al., 2015). The *C. elegans* germline was injected with a plasmid that serves as a template to transcribe a guide RNA (gRNA) that targets the C-terminus-encoding sequence of *ccb-1.* The gRNA binding sequence (AAGAGGTACGTACAGGTACT) was inserted into the vector pDD162 (from Addgene) using the NEB Q5 Site-Directed Mutagenesis Kit. A second plasmid (pIA03) was co-injected in the germline to serve as a repair template, which includes the sequences for fluorescent protein mKate2, PSD, and a self-excising cassette containing a dominant visible marker for a rolling phenotype *(sqt-1(d)),* a heat shock-inducible Cre recombinase *(hs::Cre),* and hygromycin resistance gene *(hygR).* The progeny of injected animals was selected for hygromycin resistance and rolling phenotype. Animals were then heat shocked (34°C for 4 hours) to excise the cassette, thus ending up with the *ccb-1::mKate2::PSD* sequence only. Successful knock-in was verified by PCR (forward primer: GGGGGTACGCACATTACTG, reverse primer GGGGATGTCACAAAACGGGCTG) and Sanger sequencing (primer: GCAGCAACCACAACAGCAACAAC).

### Behavioral assays

Transgenic worms were cultivated in the dark at 20°C on nematode growth medium (NGM) with OP50-1 bacteria (Brenner, 1974) without or with all-*trans* retinal (ATR; Liewald et al., 2008). Dishes containing ATR were prepared by spreading 320 μl of OP50-1 culture, mixed with 0.64 μl of 100 mM ATR stock (dissolved in ethanol) onto 5.5-cm dishes containing 8.2 ml of NGM. About 18 h prior to the experiments, L4 larvae, grown on ATR plates, were placed on fresh ATR plates. Worms were illuminated with blue light (contraction-assay: 1.4 mW/mm^2^, swimming-assay: 0.53 mW/mm^2^) from a 50 W HBO mercury lamp, filtered through a GFP excitation filter (450–490 nm), on 5.5 cm diameter dishes, under a 10× objective in a Zeiss Axiovert 40 microscope (Zeiss, Germany). The duration of illumination was defined by a computer-controlled shutter (Sutter Instruments). Worms were filmed with a Powershot G9 digital camera (Canon, Japan) at 640 × 480 resolution with 30 fps. The body length was determined as previously described (Erbguth et al., 2012). The length values were normalized to the averaged values measured before illumination (0–5 s; normalization was carried out for each animal). To exclude measurement errors (i.e. when the animals body touches itself), all values below 80% were excluded and the length-profiles were averaged for each strain. The experiments were repeated on 2–3 different days (worms were picked from different populations); the final graphs show the average of all individual animals.

For analyzing swimming behavior, worms were placed into 96-well plates containing 100 μl NGM and 100 μl of M9 buffer per well. For analysis of the effect of CCB-1 photodegradation, animals were illuminated for 1h (470 nm, 400 μW, 20°C). Worms were left in M9 buffer for 15 min before the swimming-assay, then they were filmed for 60 s. The swimming cycles (the animals’ body bends twice per cycle), were counted manually. The swimming assays were repeated on three different days, with 9–10 worms/group, picked from different populations.

Crawling speed was measured by the multi-worm tracker, as described previously (Swierczek et al., 2011).

### Pharmacological (Aldicarb and Levamisole) Assays

To assay aldicarb sensitivity, 2 mM aldicarb dishes were prepared (Mahoney et al., 2006). After transferring the animals (14–15 young adults/trial, in total three trials on three different days, with worms picked from different populations) to the dishes, they were scored every 30 min by three gentle touches with a hair pick. The assays were performed blinded and on the same day with the same batch of aldicarb dishes. For analysis of the effect of CCB-1 photodegradation, animals were illuminated for 1h (470 nm, 100 μW, 1h, 20°C), and then incubated on 1 mM aldicarb plates. The data for the 1.5 and 2 h time points were averaged.

For levamisole assays, we used an agonist of the levamisole-sensitive nAChR of the neuromuscular junction, the racemic mixture tetramisole, at 2 mM on NGM plates (corresponding to 1 mM levamisole). Animals were incubated on these plates as described for the aldicarb assay, and assessed for paralysis every 30 min.

### Electrophysiology

Electrophysiological recordings from body wall muscle cells were conducted in immobilized and dissected adult worms as described previously (Liewald et al., 2008). Animals were immobilized with Histoacryl glue (B. Braun Surgical, Spain) and a lateral incision was made to access neuromuscular junctions (NMJs) along the anterior ventral nerve cord. The basement membrane overlying body wall muscles was enzymatically removed by incubation in 0.5 mg/ml collagenase for 10 s (C5138, Sigma-Aldrich, Germany). Integrity of body wall muscle cells and nerve cord was visually examined via DIC microscopy. Recordings from body wall muscles were acquired in whole-cell patch-clamp mode at room temperature (20-22°C) using an EPC-10 amplifier equipped with Patchmaster software (HEKA, Germany). The head stage was connected to a standard HEKA pipette holder for fire-polished borosilicate pipettes (1B100F-4, Worcester Polytechnic Institute, Worcester, MA, USA) of 4-9 MΩ resistance and recordings were sampled at 3.33 kHz.

The extracellular bath solution consisted of 150 mM NaCl, 5 mM KCl, 5 mM CaCl_2_, 1 mM MgCl_2_, 10 mM glucose, 5 mM sucrose, and 15 mM HEPES (pH 7.3 with NaOH,~330 mOsm). The internal/patch pipette solution consisted of K-gluconate 115 mM, KCl 25 mM, CaCl_2_ 0.1 mM, MgCl_2_ 5 mM, BAPTA 1 mM, HEPES 10 mM, Na_2_ATP 5 mM, Na_2_GTP 0.5 mM, cAMP 0.5 mM, and cGMP 0.5 mM (pH 7.2 with KOH,~320 mOsm). With the solutions used, reversal potentials are about +20 mV for nicotinic ACh receptors (nAChRs) and −30 mV for GABA_A_ receptors (Maro et al., 2015). For most experiments, recordings were conducted at a holding potential of −60 mV where nAChR-related currents (EPSCs) and GABA_A_ receptor-related currents (IPSCs) both display as inward currents. At a holding potential of −10 mV (**Fig. S6A**), however, EPSCs and IPSCs are recorded as inward and outward currents, respectively.

Light activation was performed using an LED lamp (KSL-70, Rapp OptoElectronic, Hamburg, Germany; 470 nm, 8 mW/mm^2^) and controlled by the Patchmaster software. Subsequent analysis and graphing was performed using Patchmaster, and Origin (Originlabs). Analysis of mPSCs was conducted with MiniAnalysis (Synaptosoft, Decatur, GA, USA, version 6.0.7) to acquire parameters such as mPSC frequency, peak amplitude, time to peak (time from start of light stimulus until peak maximum), rise time (time from start of peak until peak maximum), decay time, and area under the curve. For the analysis of quantal content, the area under the curve of photo-evoked currents was divided by the mean area under the curve of respective mPSCs of the same animal measured before photostimulation.

### Fluorescence Microscopy

For analyzing the fluorescence level of GFP::UNC-2 and CCB-1::mKate2::PSD in the nerve ring, animals were transferred onto 10% agarose pads in M9 buffer (K2PO4 20 mM; Na_2_HPO_4_ 40 mM; NaCl 80 mM; MgSO4 1 mM) and immobilized with 1 μl 20 mM tetramisole hydrochloride. For the widefield fluorescence imaging of these animals, an Axiovert 200 or an Observer Z1 microscope (Zeiss, Germany) was used, equipped with a 40x/1.3 Zeiss oil objective, a 460 nm LED (Prior Scientific), a GFP filter cube (ex. 470/40 nm, em. 520/35 nm) and RFP filter cube (ex. 580/23 nm, em. 625/15). The nerve ring region was located using white transmission light. The fluorescence excitation light was turned on and a fluorescence image was captured using an EMCCD camera (Photometrics Evolve 512 Delta), gain (GFP: 68 (ZM9583), 4 (ZX2321); RFP: 20 (ZX2321, ZX2271), exposure time 50 ms (ZM9583) or 100 ms (ZX2321, ZX2271), controlled using Micro Manager software (Edelstein et al., 2014). Images were analysed using FIJI software (Schindelin et al., 2012). For quantification, a ROI was placed around the nerve ring and mean fluorescence intensity was measured. For background correction, a background value was measured in the worm’s head close to the nerve ring.

### Confocal Microscopy

The expression patterns of *rimb-1p::GFP* and *unc-17p::mCherry* (strain ZX2474) were analyzed using a spinning disc confocal microscope (Cell Observer SD, Zeiss, Germany), equipped with a 40x air objective (LD Plan-NEOFLUAR 40x/0.6 PH2 Kor ∞/0 1,5; Zeiss, Germany), an LED-based illumination system (excitation wavelengths 488 nm and 561 nm) and a double band pass emission filter (DBP 527/54 + 645/60, Zeiss). Micrographs were taken with an EMCCD camera (Rolera EM-C2, Teledyne QImaging). Animals were transferred onto 10% agarose pads in M9 buffer and immobilized with Polybead polystyrene 0.1 mm microspheres (Polysciences Inc., Warrington, PA, USA).

For analyzing the synaptic puncta intensity and size of GFP::UNC-2 in the nerve ring, animals were immobilized as described above (10% agarose pads in M9; 1 μl 20 mM tetramisole hydrochloride). In this case the spinning disc confocal microscope (Cell Observer SD, Zeiss, Germany), was equipped with a 63 x oil immersion objective (63 x/ 1.4 Oil DIC Plan apochromat), LED illumination (488 nm) and a 520/35 nm emission filter. Gain (150) and exposure time (1000 ms) were controlled by ZEN 2 blue software (Zeiss). Images were obtained with a Rolera EM-C2 EMCCD camera (Teledyne QImaging). After locating the nerve ring using only white light, z-stacks were acquired using a piezo focused stage (NanoScan Z, Prior Scientific). 40 images (0.5 μm spacing) were taken per animal, which were then z-projected (standard deviation) and analysed using FIJI software.

### Electron Microscopy

Transgenic L4 worms were transferred from regular NGM dishes to freshly seeded E. coli OP50 −/+ (0.1 mM) ATR dishes 1 to 2 days before high-pressure freezing (HPF). Young adult animals were used for HPF fixation, based on methods previously described (Rostaing et al., 2004; Weimer, 2006; Kittelmann et al., 2013). Briefly, about 10–40 worms were loaded into a 100 μm deep aluminum planchette (Microscopy Services) filled with *E. coli* OP50 −/+ ATR, covered with a 0.16 mm sapphire disc and a 0.4 mm spacer ring (Engineering Office M. Wohlwend GmbH) for subsequent photostimulation. To prevent preactivation, all manipulations were done under red light. For light stimulation experiments, worms were continuously illuminated with a laser (~20 mW/mm^2^) for 30 s, followed by HPF at −180°C under 2100 bar pressure in a Bal-Tec HPM010 HPF machine. A ~5 s period of manual insertion of the sample after photostimulation is required with this high pressure freezer.

After freezing, specimens were transferred under liquid nitrogen into a Reichert AFS machine (Leica) for freeze substitution. Tannic acid (0.1% in dry acetone) fixative was used to incubate samples at −90°C for 100 h. Then, a process of washing was performed, followed by an incubation of 2% OsO4 for 39.5 h (in dry acetone) while slowly increasing the temperature up to room temperature. Afterwards, the process of embedding in Epoxy resin (Agar Scientific, AGAR 100 Premix kit hard) was executed with increasing concentration from 50% to 100% at room temperature and 100% at 60°C over 48 h.

For electron micrographs, cross sections were cut at a thickness of 40 nm, transferred to formvar-covered copper slot grids and counterstained in 2.5% aqueous uranyl acetate for 4 min, followed by washing with distilled water. Then, grids were carried onto Reynolds lead citrate solution for 2 min in a CO2-free chamber and subsequently washed in distilled water again. Images of regions in the ventral nerve cord were taken with an Erlangshen ES500W CCD camera (Gatan) in a Philips CM12 transmission electron microscope operated at 80 kV. Images were scored blind for each condition and tagged in ImageJ (NIH). ImageJ ROIs were stored and then quantified based on methods described previously (Steuer Costa et al., 2017). The diameters of synapses from each stimulation condition vary due to the different extent of SV exocytosis or because different synapses were sampled. Thus, each value for the number of docked SVs was normalized and represents the number of docked SVs along a membrane whose perimeter is 1548 nm in a profile; the other organelles are represented as the numbers of SVs or LVs in a typical synaptic profile of 164,100 nm^2^ (Steuer Costa et al., 2017). SV size was scored from 3 to 10 middle images of randomly selected synapses or from randomly selected single images per synapse for each mutant or stimulation protocol. 2–3 worms, and typically 10–19 synapses were analyzed per genotype and condition; 1–2 technical replicates were performed.

### Voltage imaging

For voltage imaging experiments, ATR (Sigma-Aldrich, USA) had to be supplied to the animals. Therefore, one day prior to each experiment, transgenic L4 stage worms were transferred onto NGM plates, seeded with OP50 bacterial suspension supplemented with ATR (final ATR concentration: 0.01 mM). For imaging, animals were immobilized with polystyrene beads (0.1 μm diameter, at 2.5% w/v, Sigma-Aldrich) on top of 10 % agarose pads (in M9 buffer). Voltage-dependent fluorescence of Arch(D95N) was excited with a 637 nm red laser (OBIS FP 637LX, Coherent) at 1.8 W/mm^2^ and imaged at 700 nm (700/75 ET bandpass filter, integrated in Cy5 filter cube, AHF Analysentechnik), while ChR2(H134R) was stimulated by a monochromator (Polychrome V) at 300 μW/mm^2^. Imaging was performed on an inverted microscope (Zeiss Axio Observer Z1), equipped with a 40x oil immersion objective (Zeiss EC Plan-NEOFLUAR 40x/ N.A. 1.3, Oil DIC ∞ / 0.17), a laser beam splitter (HC BS R594 lambda/2 PV flat, AHF Analysentechnik), a galilean beam expander (BE02-05-A, Thorlabs) and an EMCCD Camera (Evolve 512 Delta, Photometrics).

### Statistical Analysis

All quantitative data were reported as mean ± s.e.m., and n indicated the number of animals, N the number of replicates. Significance between data sets after two-tailed Student’s t-test or after one-way or two-way ANOVA, with Bonferroni’s multiple comparison test, Fisher test, or Tukey’s post-hoc test, is given as p-value. If data was not distributed normally, we used Newman-Keuls test, or Kruskall-Wallis test with Dunn’s multiple comparisons. For paralysis assays, we used the log rank test (with Bonferroni correction) to compare datasets across the duration of the experiment. The respective statistics used are indicated for each experiment in the figure legends. * signifies p<0.05, **p<0.01, and ***p<0.001. Data was analyzed and plotted in GraphPad Prism (GraphPad Software, Inc., La Jolla, CA, USA, version 8.02), Microsoft Excel 2016, or in OriginPro 2020b (OriginLab Corporation, Northampton, USA).

## Acknowledgements

We thank members of the Gottschalk lab for critically reading the manuscript. We acknowledge the *Caenorhabditis* Genetic Center, which is funded by NIH Office of Research Infrastructure Programs (P40 OD010440), and the National Bioresource project, nematode *C. elegans*, for strains. We further thank Mei Zhen, Erik Jorgensen and Bob Goldstein for additional strains and transgenes, and Daniel J. Dickinson for advice on CRISPR gene editing. We are indebted to Franziska Baumbach, Hans-Werner Müller, Annabel Klaus, Dennis Vettkötter, Jens Gruber, Heinz Schewe, and Mona Höret for expert technical assistance. This work was supported by the Deutsche Forschungsgemeinschaft (DFG), grants CRC1080-B02 and CRC807-P11, to A.G., and by Goethe University Frankfurt. Ivan C. Alcantara was a Fulbright scholar.

## Author contributions

Experiments were conceived, designed, and analyzed by B.J., J.F.L., S.-c. Y., S.U., I.C.A., A.C.F.B., M.W.S., and A.G. Experiments were conducted by B.J., J.F.L., S.-c. Y., S.U., I.C.A., A.C.F.B., M.W.S., and J.S. The manuscript was written by B.J., J.F.L., and A.G. Funding was acquired by A.G.

## Supplemental Figures and Legends

**Figure S1:**
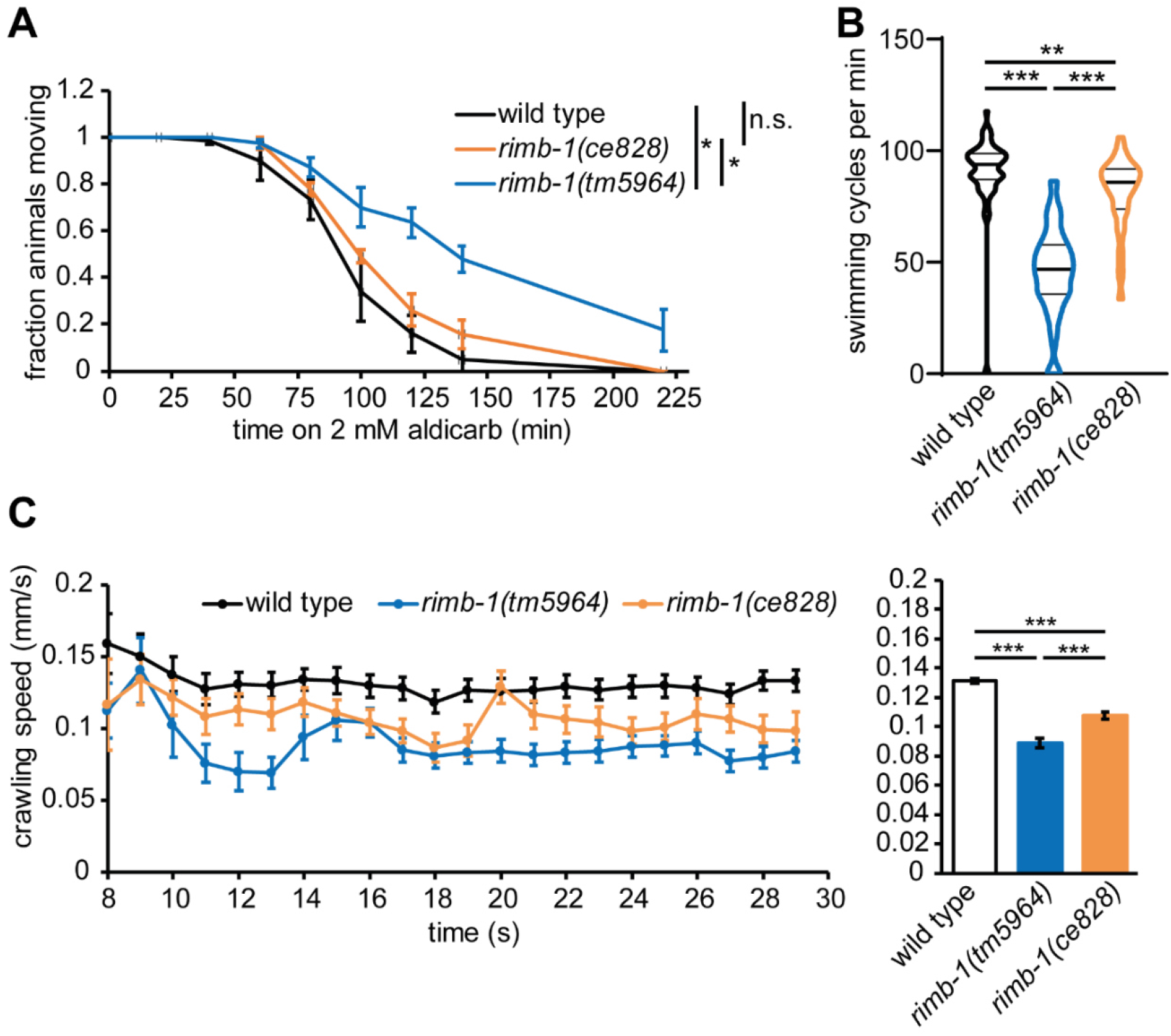
Comparing two *rimb-1* alleles. *rimb-1* mutants *tm5964* and *ce828* were compared by **A)** resistance to aldicarb, testing for cholinergic transmission (3 replicates, n=15 animals each; Log rank test, Bonferroni correction), **B)** swimming and **C)** crawling locomotion. In A and C, mean ± s.e.m. are shown, in B, median and 25/75 quartiles (thick and thin lines), min to max. One-way ANOVA with Tukey test. Statistical significance given as *p<0.05; **p<0.01; ***p<0.001.

**Figure S2:**
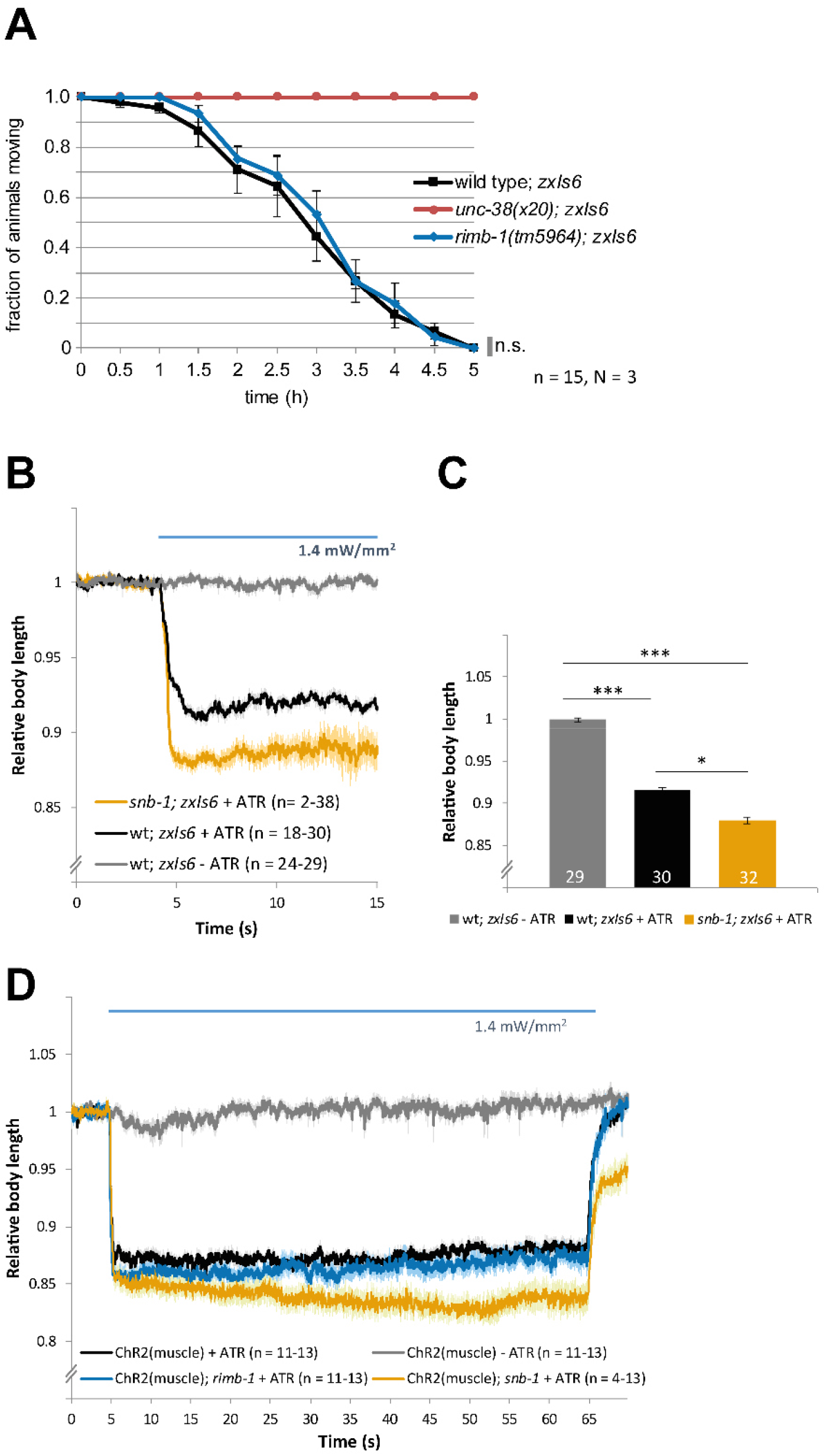
*rimb-1* mutants have no post-synaptic defect in muscular nAChRs, and normal muscle excitability. **A)** Levamisole-induced paralysis (1 mM, targeting muscular nAChRs), assessed over time, in three replicates, for the indicated number of animals of the indicated genotypes. Mean ± s.e.m. Log rank test. **B)** ChR2 expressed in cholinergic neurons, photoinduced muscle contraction (photostimulus indicated by blue bar). Shown is the mean (± s.e.m.) relative body length, n = number of animals. Synaptobrevin *snb-1(md247*) mutants show increased muscle contraction, despite reduced ACh release, due to compensatory excitability increase in post-synaptic muscle (Liewald et al., 2008). **C)** Statistical analysis of the data shown in B, one-way ANOVA, Bonferroni-corrected. **D)** Body contraction induced by ChR2 expressed in muscle, demonstrating increased contractions in *snb-1* mutants, but normal excitability in *rimb-1* mutants. Statistical significance given as ***p<0.001.

**Figure S3:**
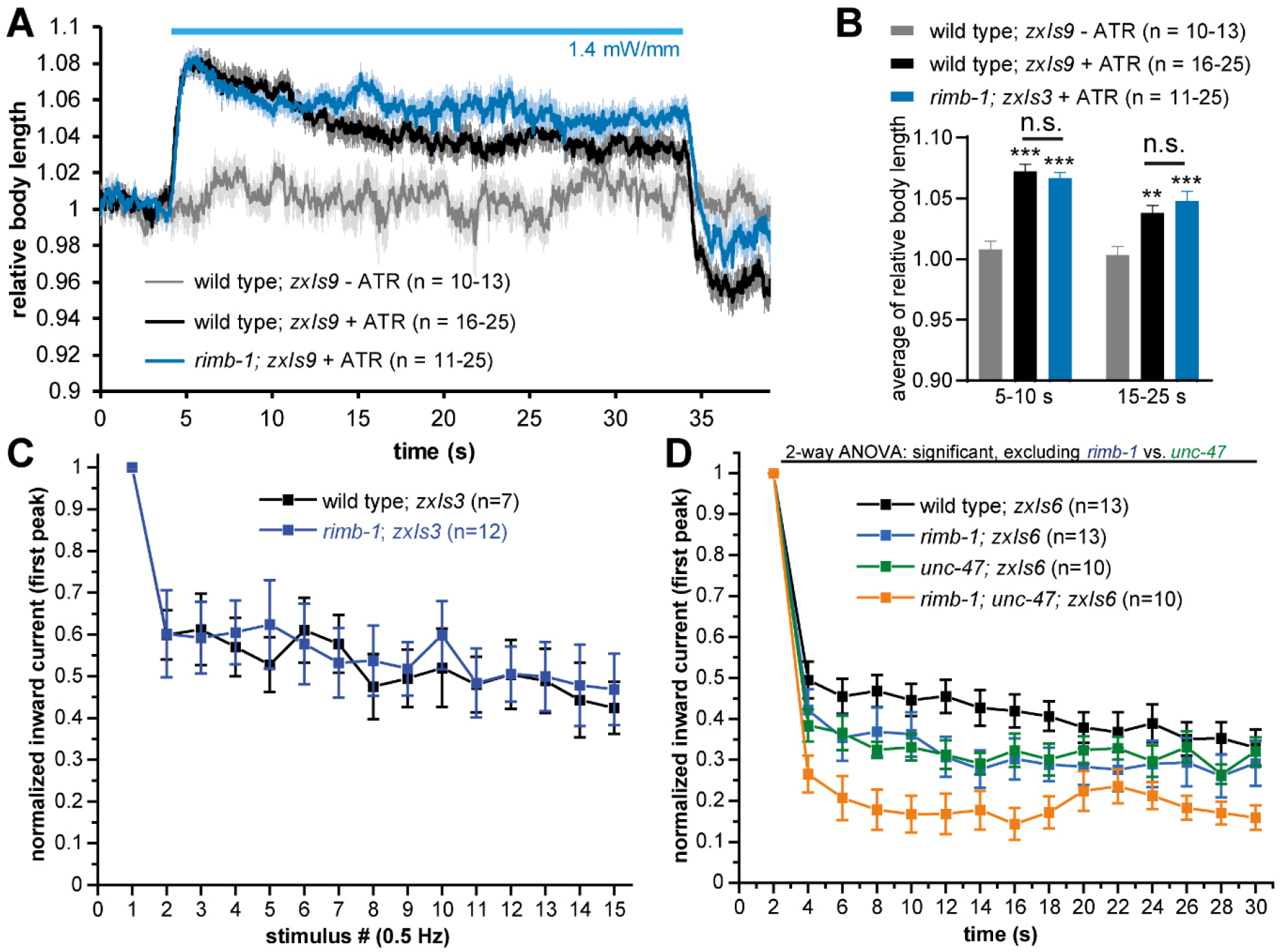
GABAergic transmission at the NMJ in *rimb-1* mutants. **A)** Photostimulation of GABAergic neurons (transgene *zxIs9*) causes body elongation. Genotypes and number of animals tested are indicated. **B)** Group data of the experiments shown in A, for the time period 1-5 s of the photostimulus; one-way ANOVA with Bonferroni correction. Statistical significance given as ***p<0.001. **C)** Photoevoked IPSCs for 15 consecutive stimuli, normalized to the first peak; means ± s.e.m. Related to **Fig. 5D**. No progressive change in the normalized amplitude is apparent. **D)** Indirectly probing the GABAergic NMJ using cholinergic stimulation of GABAergic neurons, comparing wild type, *rimb-1* and *unc-47* vGAT mutants, normalized to the first peak. Related to **Fig. 6E**. Two-way ANOVA with Fisher test.

**Figure S4:**
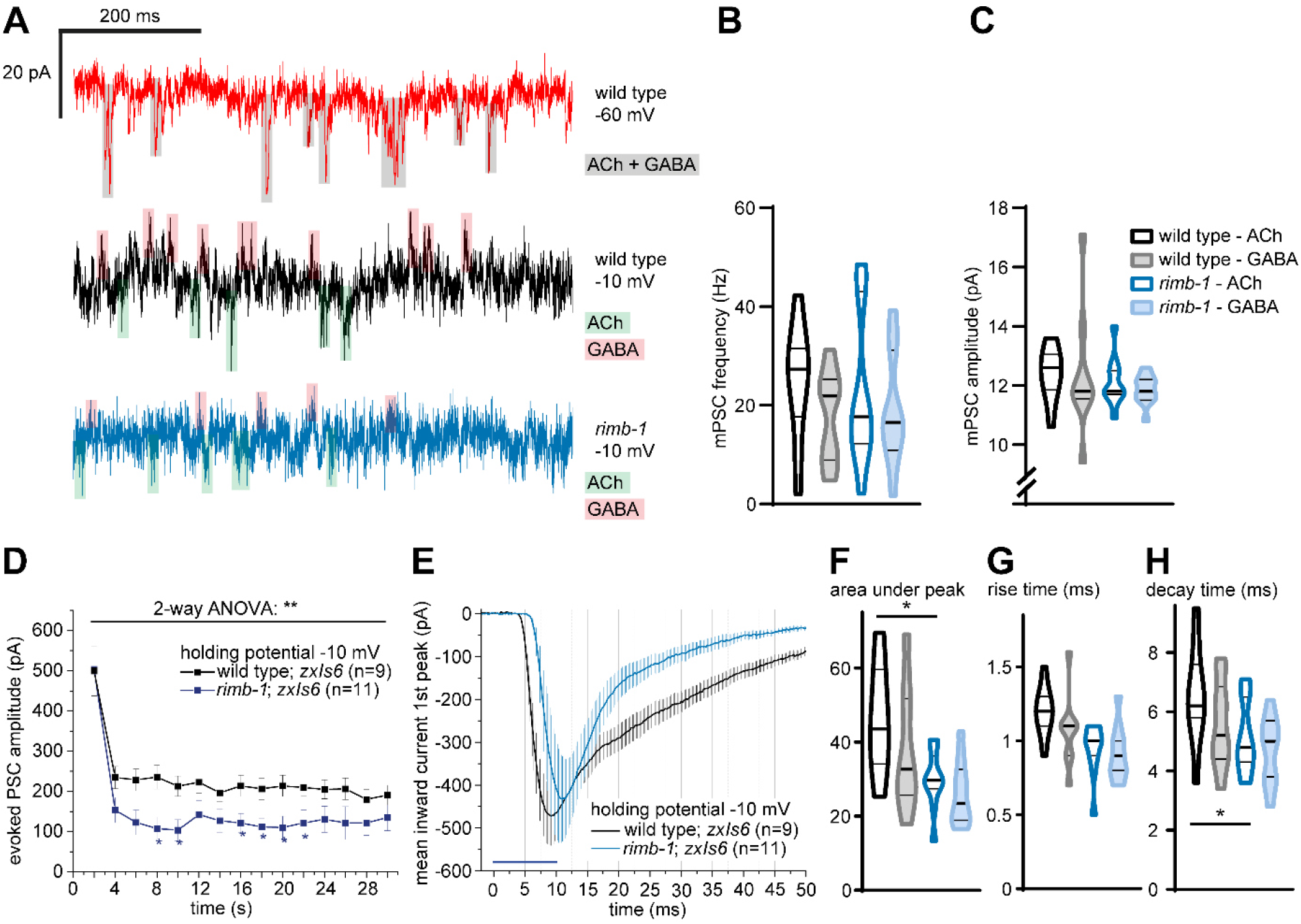
*rimb-1* defects in evoked NMJ transmission at −10 mV. **A)** mPSCs measured at different holding potentials (−60 mV or −10 mV), discerning cholinergic (green shade) and GABAergic (red shade) currents. Genotypes as noted. **B)** Frequency and **C)** amplitude, of cholinergic and GABAergic mPSCs, in wild type and *rimb-1(tm5964*) mutants. **D)** Mean (± s.e.m.) ePSCs for 15 consecutive cholinergic photo stimuli, measured at −10 mV. n = number of animals. **E)** Time course of first ePSC (mean ± s.e.m.), measured at −10 mV. Analysis of mPSC parameters **F)** area under peak, **G)** rise and **H)** decay time. One-way ANOVA with Tukey test. Data in B, C, F-H shown as median and 25/75 quartiles (thick and thin lines), min to max. Statistical significance in D, F given as *p<0.05; **p<0.01.

**Figure S5:**
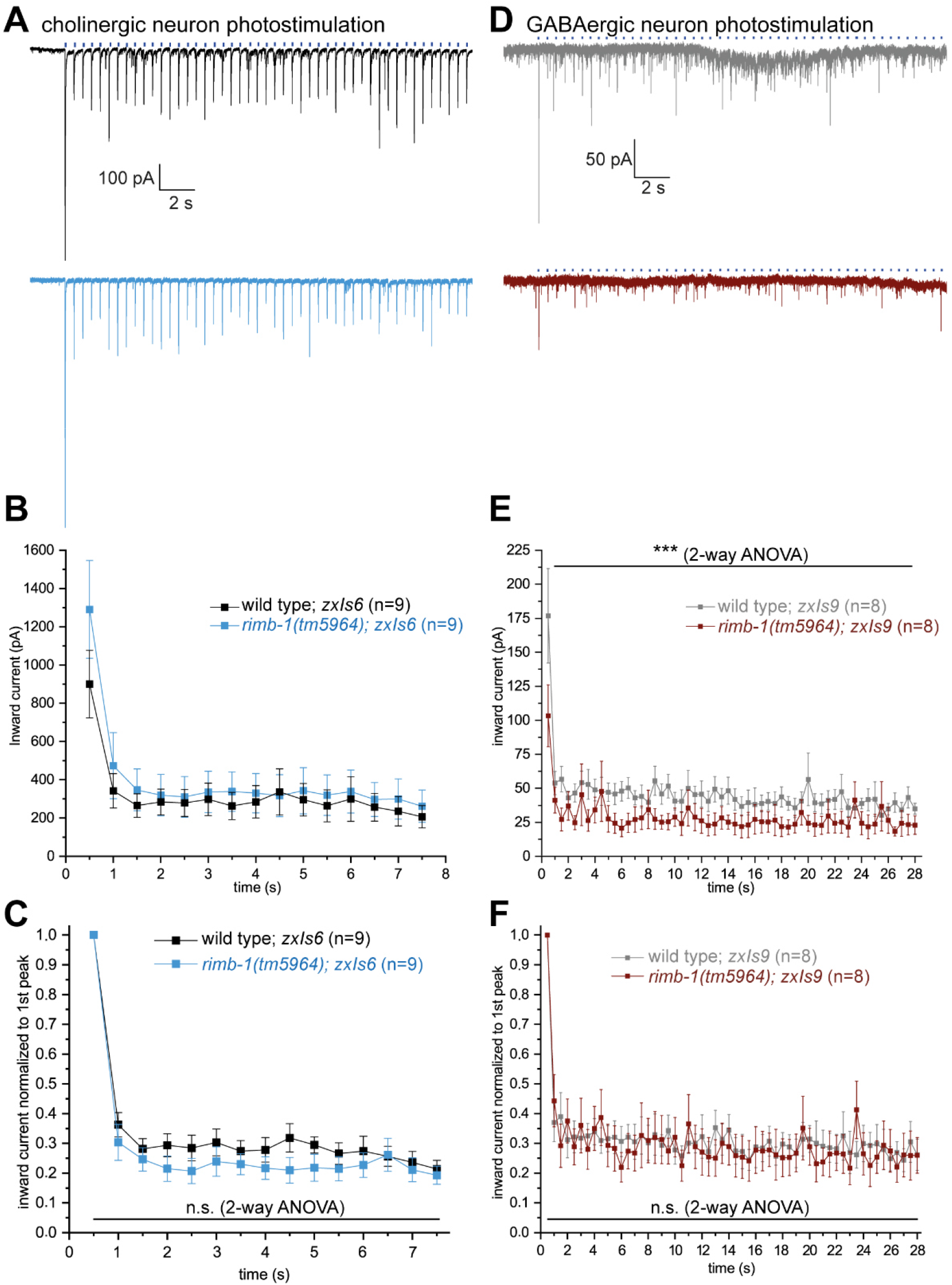
Repeated 2 Hz stimulation of cholinergic and GABAergic synapses reveals no effect on depression or facilitation. **A-C)** Cholinergic (transgene *zxIs6*) and **D-F)** GABAergic (transgene *zxIs9*) photostimulation, 2 Hz, 10 ms stimuli. Original records, mean (± s.e.m.) ePSCs, without or with normalization to the first peak. **D-F)** Cholinergic photostimulation, 2 Hz, 10 ms stimuli. Original records, mean (± s.e.m.) ePSCs, without or with normalization to the first peak.

**Figure S6:**
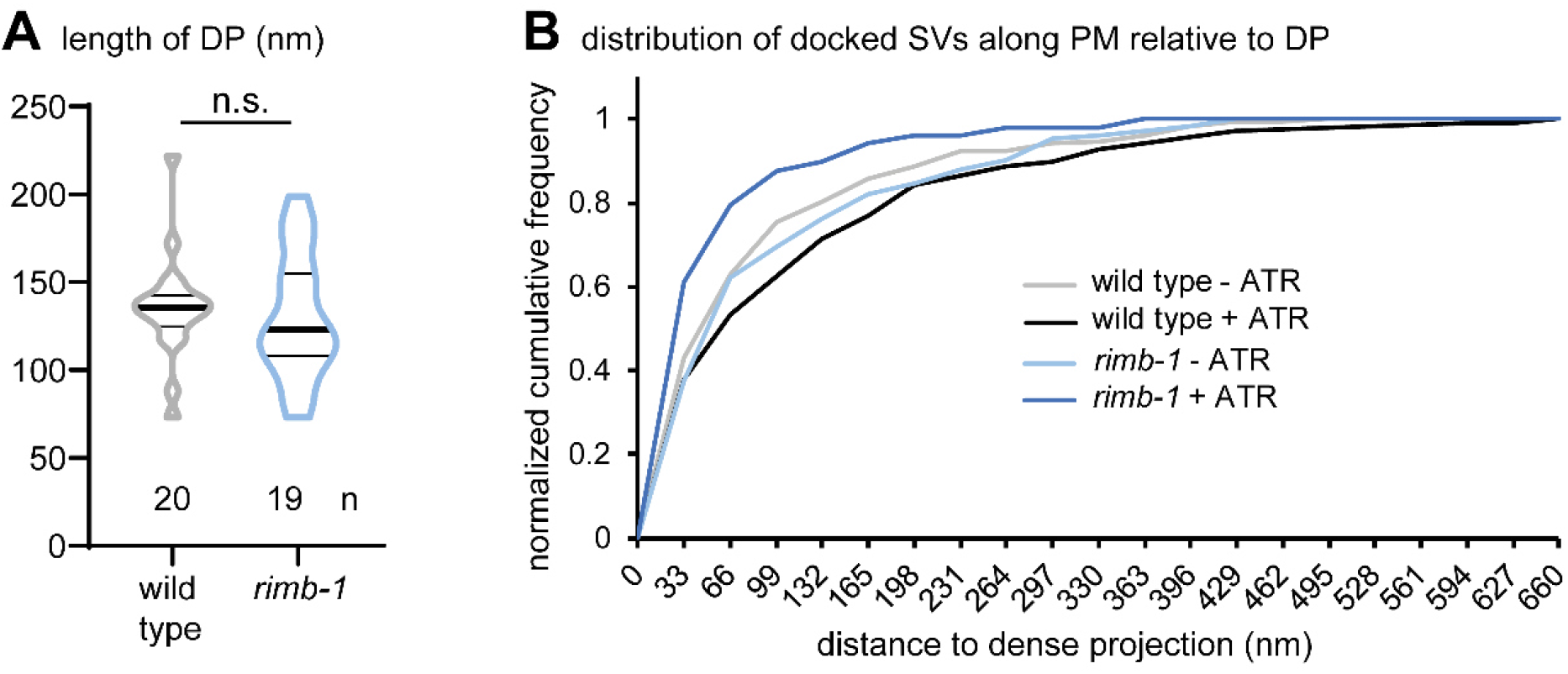
Ultrastructural parameters of *rimb-1* mutant synapses. **A)** Length of the dense projection was not altered in *rimb-1* synapses. **B)** Cumulative distribution of docked SVs in illuminated wild type vs. *rimb-1* mutant synapses, with or without ATR. Docked SVs in silent synapses are not differently distributed in *rimb-1* mutants, but fuse more distal to the DP. Related to **Fig. 5F**.

**Figure S7:**
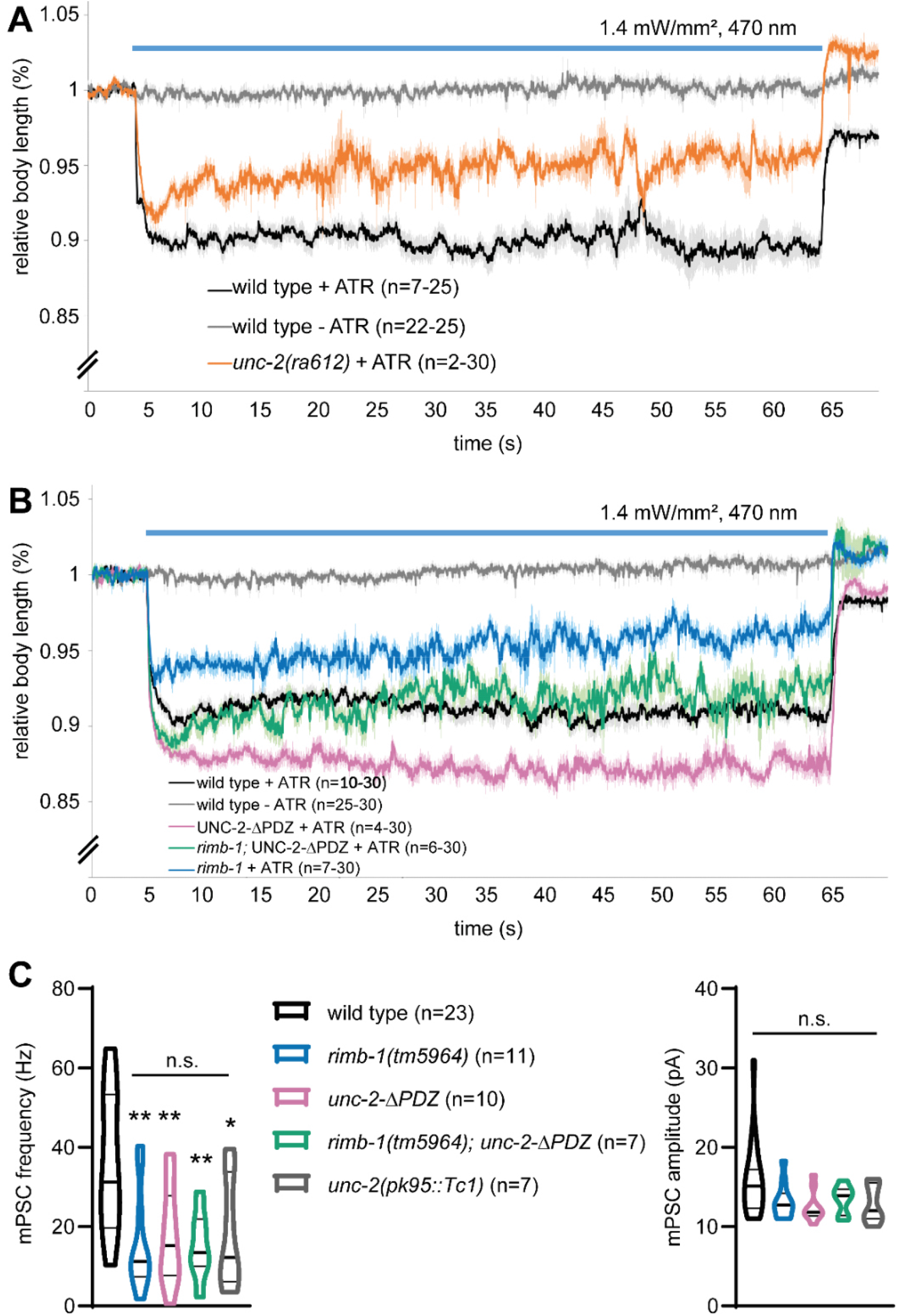
Differential effects of untethering and deleting CaV2/UNC-2 VGCCs on induced behavior and basal synaptic transmission. **A, B)** Photoinduced body contraction (transgene *zxIs6*) in *unc-2* deletion mutants, as well as *rimb-1,* UNC-2-ΔPDZ and *rimb-1;* UNC-2-ΔPDZ double mutants, compared to wild type controls. Mean ± s.e.m., light stimulus indicated by blue bar; n= number of animals analyzed. **C)** Spontaneous synaptic transmission, mPSC frequency and amplitude, compared in the indicated genotypes. n = number of animals, data shown as median and 25/75 quartiles (thick and thin lines), min to max. One-way ANOVA with Tukey test, *p<0.05, **p<0.01.

